# Properties and proximity proteomics of synaptopodin provide insight into the molecular organization of the spine apparatus of dendritic spines

**DOI:** 10.1101/2021.12.30.474557

**Authors:** Hanieh Falahati, Yumei Wu, Vanessa Feuerer, Pietro De Camilli

## Abstract

The spine apparatus is a specialization of the neuronal ER in dendritic spines consisting of stacks of interconnected cisterns separated by a dense matrix. Synaptopodin, a specific actin binding protein of the spine apparatus, is essential for its formation, but the underlying mechanisms remain unknown. We show that synaptopodin, when expressed in fibroblasts, forms actin-rich structures with connections to the ER, and that an ER-tethered synaptopodin assembles into liquid condensates. We also identified protein neighbors of synaptopodin in spines by *in vivo* proximity biotinylation. We validated a small subset of such proteins and showed that they co-assemble with synaptopodin in living cells. One of them is Pdlim7, an actin binding protein not previously identified in spines, and we show its precise colocalization with synaptopodin. We suggest that the matrix of the spine apparatus has the property of a liquid protein condensate generated by a multiplicity of low affinity interactions.

**Graphical abstract:** 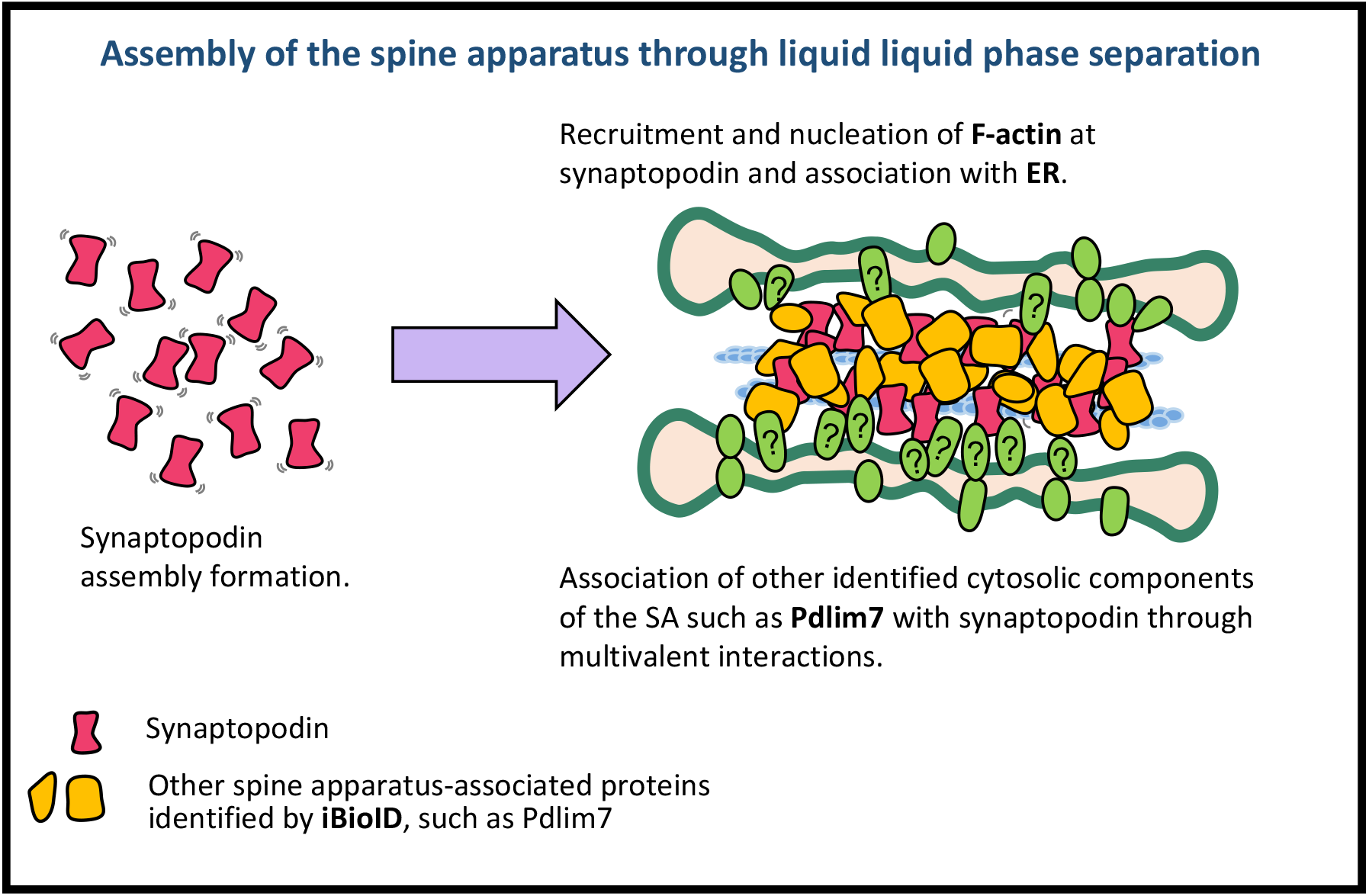

## INTRODUCTION

The neuronal ER is an intricate continuous network of membrane tubules and cisterns that runs throughout neuronal processes with region-specific specializations. One such specialization of the smooth ER is the spine apparatus (SA) that is located in a subset of dendritic spines. The SA consists of stacks of flat cisterns that are connected by an unknown dense matrix and are continuous with each other and with the ER of dendritic shaft (Gray, 1959; Spacek and Harris, 1997; Wu et al., 2017) (see also Figures 1A and S1A). Morphological changes in the SA have been reported after long-term potentiation (Chirillo et al., 2019), and also in a variety of human disorders including several neurodegenerative conditions (Baloyannis et al., 2007; Castejòn et al., 1995; Fiala et al., 2002; Špaček, 1987; Villalba and Smith, 2011). While the first observation of the SA by EM was reported in 1951 by Gray (Gray, 1959), our understanding of this organelle remains fairly limited. Its molecular characterization has proven to be challenging due to the difficulty of its biochemical isolation and its absence in organisms suitable for genetic screens.

**Figure 1.**
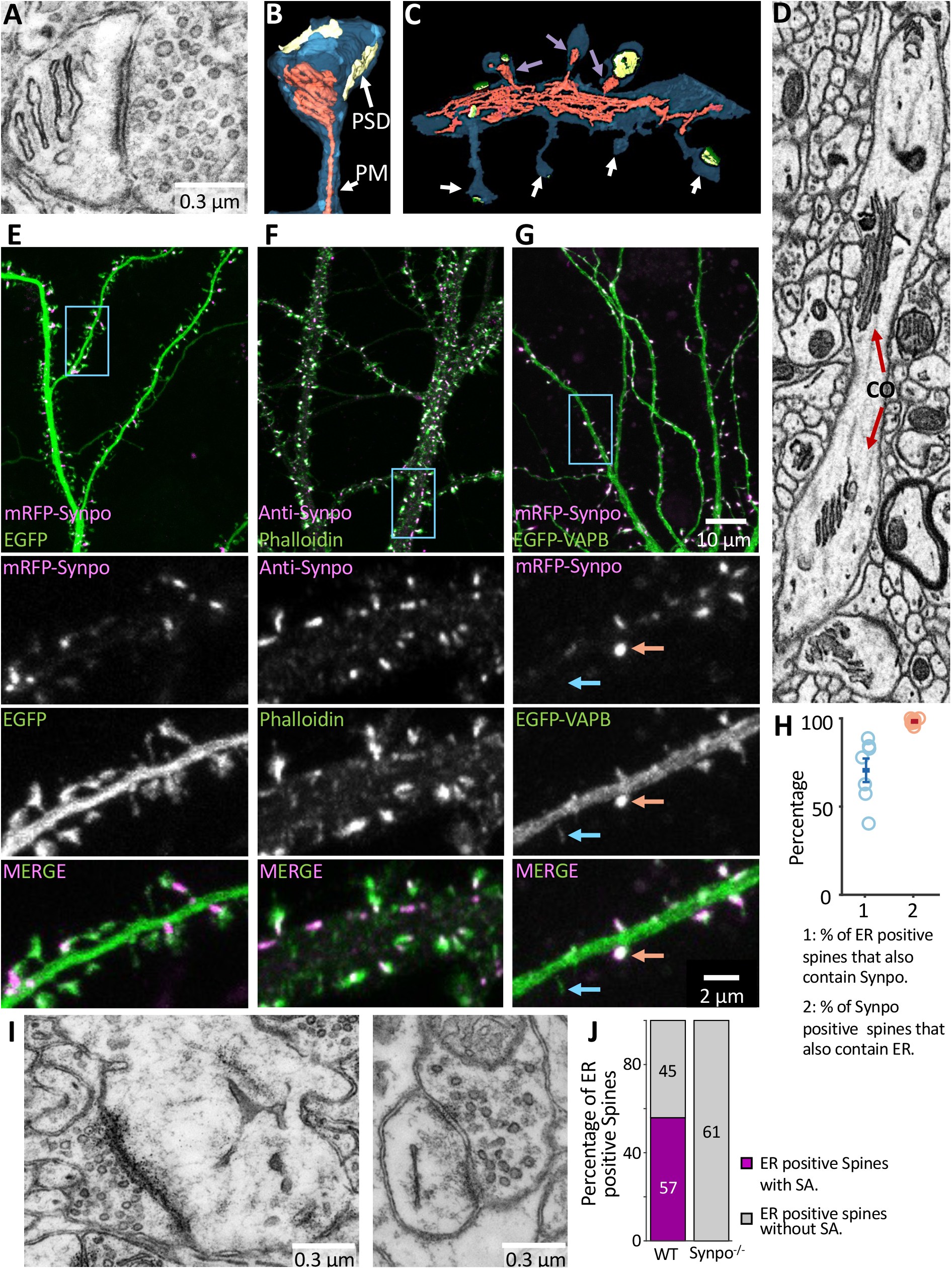
The localization of synaptopodin (indicated as Synpo in all figures) in cultured hippocampal neurons overlaps with the localization of the spine apparatus (SA) in dendritic spines of cortical slices. **A**. SA as visualized by transmission EM. **B**. and **C**. SA and ER reconstructed by a semi-automated algorithm from 3D volumes acquired by FIB-SEM. A SA is shown in B, while C shows a portion of a dendritic shaft with spines containing (magenta arrows) and not containing (white arrows) a SA. The plasma membrane is shown in blue, the ER in red, and the postsynaptic density (PSD) in yellow. **D** Cisternal organelle (CO), as observed in a FIB-SEM optical section, at an axonal initial segment (AIS). Note stacks of ER cisterns similar to those characteristics of the SA. **E**. mRFP-synaptopodin coexpressed with cytosolic EGFP as a marker of the entire dendritic volume. Note in the zoomed-in views of the region enclosed by a rectangle (bottom three fields) that synaptopodin is concentrated near the spine neck, where the SA is localized. **F**. Localization by immunofluorescence of endogenous synaptopodin showing strong overlap with a pool of F-actin labeled by phalloidin-Alexa 488. Also, in this sample the magnified views (bottom three fields) show enrichment of synaptopodin, relative to actin, at the spine neck. **G**. mRFP-synaptopodin coexpressed with an ER marker, EGFP-VAPB, showing colocalization of the two proteins. In the zoomed-in views (bottom three fields), red arrows show a spine positive for both ER marker and synaptopodin, and the blue arrows show a spine with ER marker but lacking synaptopodin. **H**. Percentage of ER positive spines that also contain synaptopodin, and percentage of synaptopodin positive spines which contain ER quantified in cultured hippocampal neurons expressing mRFP-synaptopodin and the ER markers EGFP-VAPB or EGFP-Sec61ß. Each datapoint represents at least 99 spines from a single neuron. **I**. Spines of a synaptopodin KO mouse that lack the SA but contain ER (**C**). **J** Quantification of the number of spines where ER was visible in the plane of section with, or without a SA in the brain of WT versus synaptopodin mutant mice (n = > 800 spines per genotype).

The only known protein enriched at the SA and required for its formation is synaptopodin, a protein without transmembrane regions localized in the cytosolic space (Mundel et al., 1997). Neuronal synaptopodin specifically localizes to dendritic spines and to the axonal initial segment where another specialization of the ER similar to the spine apparatus (stack of flattened cisterns) called the cisternal organelle, is present (Bas Orth et al., 2007; Deller et al., 2000; Sánchez-Ponce et al., 2012; Vlachos, 2012). Lack of synaptopodin in synaptopodin KO mice correlates with lack of SA and of the cisternal organelle, as well as with a reduction in Hebbian plasticity and spatial memory (Bas Orth et al., 2007; Deller et al., 2003, 2007; Vlachos et al., 2009; Yap et al., 2020a). Synaptopodin binds to and bundles actin (Asanuma et al., 2005) and interacts with several actin binding proteins, such as α-actinin (Asanuma et al., 2005; Kremerskothen et al., 2005; Sánchez-Ponce et al., 2012). A longer isoform of synaptopodin is expressed in the foot processes of podocytes, where it functions as a regulator of the actin cytoskeleton (Asanuma et al., 2006; Patrie et al., 2002). How these interactions contribute to the formation of the spine apparatus remain unknown.

The goal of this work was to gain new insight into the molecular properties of the SA. To this aim we used synaptopodin as a starting point for our analysis. We studied its assembly in cells overexpressing it with or without some of its binding partners that we have identified by an in vivo proximity biotinylation approach. Our results reveal the property of synaptopodin to form liquid condensates that can be associated with the ER and are mediated by a multiplicity of low affinity interactions. We propose that phase separation may contribute to the formation of the SA.

## RESULTS

### Localization of synaptopodin correlates with presence of the SA in neurons

As a premise to our molecular analysis, we examined the morphology of the SA by electron microscopy. Representative electron micrographs of the SA are shown in Figures 1A and S1A. We also analyzed 3D volumes of cerebral cortex by FIB-SEM to assess average morphology and localization of the SA within spines. FIB-SEM is a technique that allows the visualization of the entire 3D volume of the spine (Wu et al., 2017; Xu et al., 2017). To this aim, we developed an algorithm to semi-automatically detect the SAs in FIB-SEM images. We defined the SA as a portion of ER with two or more closely apposed parallel flat cisterns located in dendritic spines. To detect a SA, a segment of it was manually selected in one plane, and its remaining portion was automatically tracked in the other parallel planes. Using this algorithm, we identified and examined 83 SAs in FIB-SEM images of three samples of mouse cerebral cortex reported previously (Hayworth et al., 2015) (see examples in Figures 1B-C, S1B and Video 1). Typically, the SA was located close to boundary of the spine head with the spine neck (Figures 1B and S1B-C). The ER is known to form contacts with other membranes where ions or lipids are exchanged. 83% of the identified SAs had at least one site of close appositions with the PM, while 41% of them were in contact with one or more tubulovesicular structures not connected to the ER, most likely endosomes (see Figure S1C for examples). The number of ER cisterns within SAs varied between 2 and 8, with the median number being three (Figure S1D). A dense matrix was not only present between cisterns but also in some cases at the free surface of one of the two outer cisterns of the SA (Figures 1A and S1A). As expected, analysis of FIB-SEM 3D volumes also revealed stacks of ER cisterns similar to those of the SAs, but more extended in width, at axonal initial segments where they are called cisternal organelles (Figure 1D).

Inspection of hippocampal neurons in primary cultures expressing mRFP-synaptopodin and the cytosolic marker EGFP confirmed presence of synaptopodin in spines (Deller et al., 2000) where it strongly colocalized with a pool of F-actin as detected by phalloidin (Figures 1E-F). Consistent with EM observations, mRFP-synaptopodin was concentrated close to the neck of the spine (Figure 1E, see high magnification fields) where the stable pool of actin is reported to reside, and was absent from the head where branched F-actin predominates (Figure 1F) (Honkura et al., 2008). In addition, in a subset of cultured hippocampal neurons endogenous synaptopodin localized to axonal initial segments where the cisternal organelle is observed by EM (Figure S1E) (Bas Orth et al., 2007; Peters et al., 1968). In agreement with synaptopodin being a component of the SA, mRFP-synaptopodin fluorescence closely overlapped with the fluorescence of the ER markers EGFP-VAPB or EGFP-Sec61ß (Dong et al., 2016), with 98% of the spines positive for synaptopodin being also positive for these ER markers (Figure 1G-H). In contrast, only 71% of spines positive for ER also contained synaptopodin puncta. Similar statistics were reported for organotypic slices (Holbro et al., 2009; Konietzny et al., 2019; Perez-Alvarez et al., 2020).

Synaptopodin is necessary for the stacking of SA cisterns, as previously reported by EM in brain tissue of synaptopodin KO mice (Deller et al., 2003) and as confirmed by us in these mice (Figure 1I-J). However, it is clearly not required to recruit ER to spines, as cultured hippocampal neurons of synaptopodin KO mice contained ER, as detected by expression of dsRed-KDEL in a subset of spines (Figure S1F). EM images further showed that ER cisterns can be present in these spines, although stacks do not form (Figure 1I-J) (Deller et al., 2003). These findings are consistent with a scenario in which synaptopodin is part of the dense cytoskeletal matrix that connects to each other cisterns of the SA.

### A pool of synaptopodin co-localizes with a subset of actin filaments associated with the ER

As mentioned above, the SA is a specialization of the ER located at sites enriched in stable actin filaments. Dendritic spines are very small cell compartments. To investigate the relationship between synaptopodin, actin and the ER, we examined the localization of fluorescently-tagged synaptopodin (mRFP-synaptopodin) in RPE1 and COS-7 cells that lack endogenous synaptopodin (Yanagida-Asanuma et al., 2007), where such relationship could be better assessed by fluorescence microscopy. In these cells, mRFP-synaptopodin colocalized with filamentous actin as visualized by phalloidin staining as described previously (Mundel et al., 1997), although with variable relative intensity (Figure 2A). A subset of mRFP-synaptopodin- and actin-positive elements in these cells had the appearance of stress fibers, and such localization was disrupted by 2µM Latrunculin A (LatA), a treatment that disrupts them (Figure 2B). On these fibers, synaptopodin had a discontinuous pattern of fluorescence, similar to that of α-actinin (Figure 2A), consistent with previous reports showing the interaction between synaptopodin and α-actinin (Asanuma et al., 2005; Kremerskothen et al., 2005). Accordingly, synaptopodin closely colocalized with EGFP-α-actinin-2 and was necessary for the localization of this protein to F-actin (Figure S2A-D), while it alternated with the localization of EGFP-myosin IIA (Figure S2E). Another subset of mRFP-synaptopodin positive elements were thick linear structures enriched in the central region of the cell (Figure 2A) and not sensitive to 2µM LatA (Figure 2C). A resistance to LatA was reported for synaptopodin clusters in dendritic spines (Vlachos et al., 2009).

**Figure 2.**
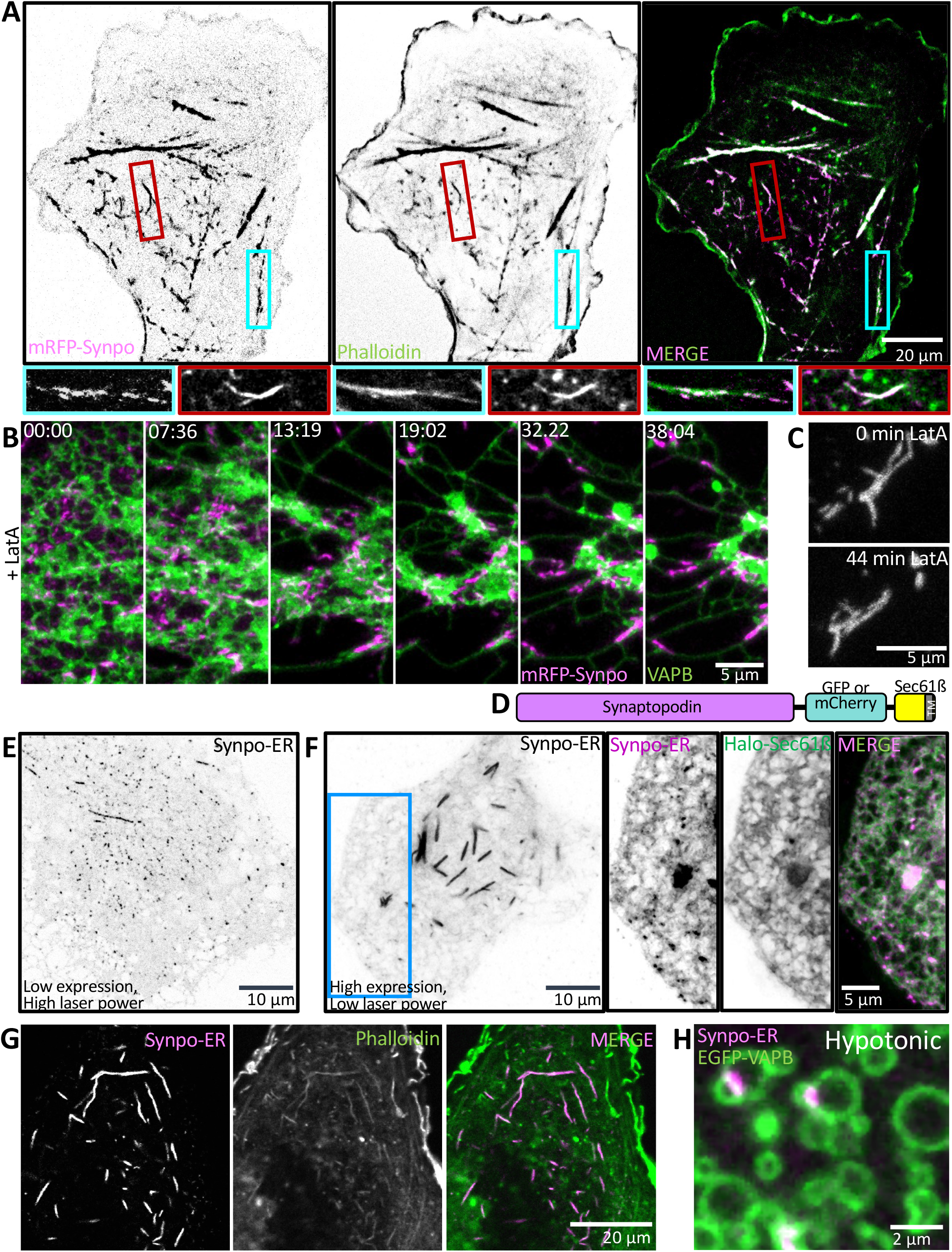
Exogeneous synaptopodin expressed in COS-7 cells localizes to actin-rich structures that can associate with ER. **A**. COS-7 cells expressing mRFP-synaptopodin showing strong overlap of the mRFP fluorescence with phalloidin staining. High magnification views of a stress fiber (cyan) and a non-stress fiber linear assembly (red) are shown at the bottom. **B-C**. Treatment of COS-7 cells expressing EGFP-synaptopodin and the ER marker mCherry-VAPB with 2µM Latrunculin A (LatA) resulted in the collapse of synaptopodin onto the ER (**B**), while the morphology of non-stress fiber synaptopodin assemblies remain unchanged (**C**). **D**. Schematic of synaptopodin-ER, a chimeric protein made by the fusion of synaptopodin to the N-terminal of the ER protein Sec61ß. E and F. Cells expressing low (**E**) and high levels (**F**) of synaptopodin-ER. At low expression levels, only small dot-like assemblies are observed (**E**), while at higher expression levels (**F**), large linear structures are also observed. The dot-like structures are not visible in F because the laser power was optimized to avoid oversaturation of the large synaptopodin assemblies. However, analysis with a different contrast and brightness of the region of **F** enclosed by a blue rectangle shows also in this cell presence of dot-like structures and their localization on the ER network labeled by the ER marker Halo-Sec61ß. **I**. The large synaptopodin-ER assemblies in COS-7 cells contain filamentous actin as revealed by phalloidin. **H**. COS-7 cells expressing synaptopodin-ER and exposed to drastic hypotonic conditions showing the localization of synaptopodin assemblies at the interface of ER vesicles.

The synaptopodin pool that was sensitive to LatA collapsed onto the ER upon LatA treatment, as shown by live microscopy. After such treatment synaptopodin clusters colocalized with, and moved in parallel to, the ER (Figure 2B; Video 2). Moreover, upon exposure to drastic hypotonic conditions, an approach previously used to examine ER contacts with other structures (Guillén-Samander et al., 2021; King et al., 2020), 90% (± 2%) of synaptopodin clusters appeared to be connected with the ER, which after such treatment reorganizes into large vesicles (Figure S2F-G, Video 3), and 65% (± 15%) of them were at contacts between ER vesicles. Such a localization supports the possibility that synaptopodin may be part of a protein network that cross-links ER cisterns.

To further investigate the interactions of synaptopodin that may underlie a role in bringing ER cisterns together, this protein was directly targeted to the ER by fusing it to the N-terminus of Sec61ß, an ER resident protein anchored to this organelle by a C-terminal transmembrane region (Zuleger et al., 2011) (Figure 2D). When expressed at low levels, this chimera, referred to henceforth as synaptopodin-ER, formed small dot-like clusters in the ER membrane (Figure 2E), while when expressed at higher levels it also resulted in the formation of large linear assemblies (Figure 2F) positive for phalloidin (Figure 2G), indicating that synaptopodin can either directly or indirectly nucleate or recruit F-actin at the ER surface. Upon exposing cells expressing synaptopodin-ER to drastic hypotonic conditions, synaptopodin-ER clusters were often found at the interface of ER vesicles (Figure 2H), further supporting the hypothesis that synaptopodin can be associated with ER membranes and bring them together.

### Synaptopodin assemblies are dynamic

In the past few years, phase separation has emerged as a mechanism contributing to the assembly of proteins into macromolecular condensates (Banani et al., 2017; Falahati and Haji-Akbari, 2019; Hyman and Brangwynne, 2011). The main drivers of this process are weak multivalent interactions of folded domains or disordered regions of proteins. Synaptopodin is predicted to be primarily a disordered protein (Figure S2H) and the features of some of the assemblies of WT synaptopodin or synaptopodin ER suggest that they have liquid-like properties. For example, fusion events between synaptopodin-ER assemblies can be observed (Figure 3A). Moreover, upon disruption of actin polymerization, WT synaptopodin associated with stress fibers collapses and relaxes into structures with smaller aspect ratio (a more compact structure) (Figure 3B). To test the hypothesis that such structures have liquid-like properties, we used fluorescence recovery after photobleaching (FRAP) on non-stress fiber synaptopodin assemblies. In COS-7 cells, the fluorescence of WT synaptopodin recovered fast, reaching 77% of its initial value within two minutes (Figure 3C-D), a time course that fits well with a single exponential curve (R^2^ = 0.99). Based on this curve we estimate that 91.3 ± 0.6% of the proteins in these assemblies are highly mobile (mobile fraction, see methods) with a recovery time constant (τ) of 1.05 ± 0.03 min (Figure 3E-F, n = 15), consistent with liquid like properties.

**Figure 3.**
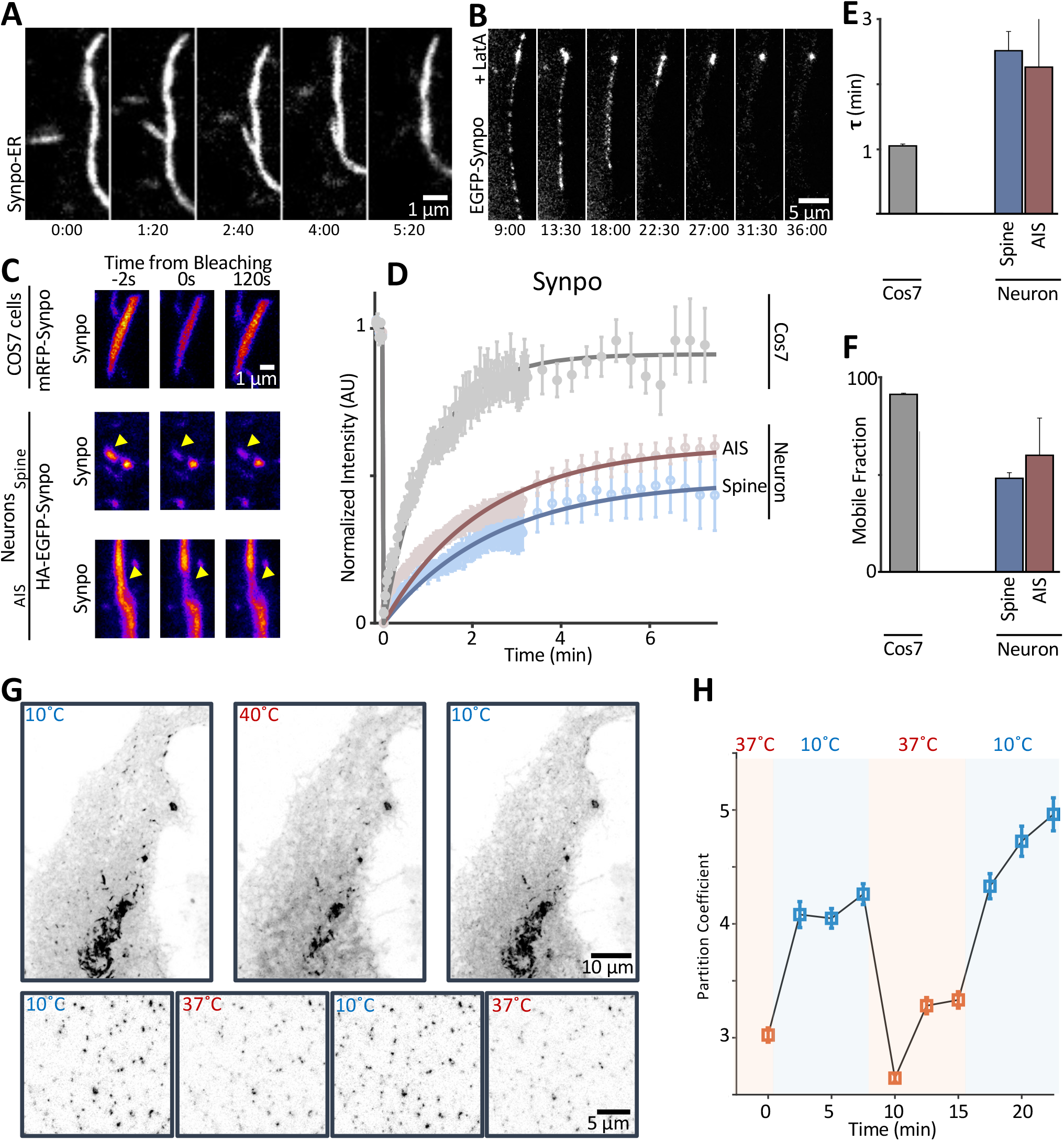
A thermodynamically-driven liquid-liquid phase separation contributes to the formation of synaptopodin assemblies. **A**. Synaptopodin-ER assemblies can fuse and coalesce. **B**. Upon disruption of F-actin by LatA treatment, EGFP-synaptopodin assemblies on stress fibers collapse into a structure with smaller aspect ratio. **C**. The dynamics of synaptopodin assemblies were assessed using fluorescence recovery after photobleaching (FRAP) in COS-7 cells or neurons. Representative images of the assemblies before bleaching (time -2s) and at 0 and 2 min after photobleaching in either COS-7 cells or neurons are shown in C. Quantifications of the recovery are shown in **D-F**. The FRAP of synaptopodin was assessed in COS-7 cells expressing mRFP-synaptopodin (n = 15) and in dendritic spines (n = 11) or axon initial segments (n = 8) of neurons (where the SA and the cisternal organelle respectively are localized) expressing HA-EGFP-synaptopodin. The lines in D show exponential fits to the fluorescent recovery curve, and each point represents mean normalized intensity ± S.E.M. All intensities are normalized to the average intensity in the first five images pre-bleaching. See methods for more details. The recovery time constant (**E**) and the mobile fraction (**F**) was calculated for each condition from the fitted single exponential curves. G. Cells expressing synaptopodin-ER in COS-7 cells were subjected to rapid temperature changes to test the role of a thermodynamically driven phase separation in the formation of synaptopodin assemblies. **H**. An approximation of the partition coefficient for synaptopodin between the assemblies and the diffuse pool is shown at different temperatures. Time interval is 2.5 ± 0.5 min.

To determine whether a thermodynamically-driven phase separation process contributes to the condensates of synaptopodin, we utilized a previously developed temperature-based approach (Falahati and Haji-Akbari, 2019; Falahati and Wieschaus, 2017; Fritsch et al., 2021) that relies on the differential response of active versus thermodynamically driven phase separation to changes in the temperature: active processes are slower at lower temperatures, due to the reduced number of favorable collisions and slower enzymatic reactions. In contrast, phase separations are often enhanced at lower temperatures because of the reduction in the entropic cost of demixing (Falahati and Haji-Akbari, 2019; Prausnitz et al., 1998). In addition, phase separation, similar to other thermodynamically driven processes, is reversible, while active processes are irreversible, and such reversibility can be tested by assessing the response to rapid changes in the temperature.

We examined the temperature-dependence of synaptopodin assembly by employing a device that allowed to change cell temperature from 37°C to 10°C within seconds (see methods for details). We chose to use synaptopodin-ER for these experiments as there seemed to be a large pool of unassembled protein for this construct, based on the fluorescence throughout the ER, compared to WT synaptopodin, where the bulk of the protein seemed to be in an assembled state (very low cytosolic fluorescence). This large pool of unassembled protein facilitates detection of a potential increase in the size of the assemblies. As can be seen in Figure 3G, a shift in the temperature from 10 to 37 °C or 40°C resulted in a decrease in the size of the assemblies. The response to changes in the temperature was reversible upon returning cells to 10°C (Figure 3G-H). This was quantified for the 10 to 37 °C shifts by calculating an approximate Partition Coefficient (PC) defined as the ratio of the concentration of synaptopodin-ER in the small assemblies relative to its concentration in the neighboring ER (Figure 3H) (See methods for details). These findings are consistent with a model in which thermodynamically driven phase separation contributes to the assembly of synaptopodin.

We next examined the dynamic properties of EGFP-synaptopodin puncta in neurons by FRAP. In dendritic spines, synaptopodin assemblies had a slower kinetics of recovery compared to COS-7 cells. 26% of the fluorescence recovered within the first two minutes of bleaching (Figure 3C-D). This recovery fits well with a single exponential curve (R^2^ = 0.95). Based on this curve we estimate that 40 ± 4% of synaptopodin in these assemblies are mobile with a time constant (τ) of 2.1 ± 0.1 min (n = 11 spines of 5 neurons) (Figure 3E-F). The slower recovery could be due, at least in part, to the confinement of the spine from the dendritic shaft. Thus, we also performed FRAP on EGFP-synaptopodin at AIS (Figure 3C-D). While the recovery at AIS was slightly faster than synaptopodin assemblies at spines, it was still slower than assemblies in COS-7 cells. This slower kinetics is likely due to the interaction of synaptopodin with other components of the SA and cisternal organelle in neurons.

### Identification of novel SA protein by proximity labeling

To identify new components of the SA that may contribute to synaptopodin in forming this structure, we used an in vivo proximity-labeling approach (Figure 4A) (Uezu et al., 2016a). We generated a fusion protein of synaptopodin and BioID2 (Kim et al., 2016) (BioID2-synaptopodin) and confirmed that this protein, when expressed in hippocampal cultured neurons, was targeted to dendritic spines by anti-BioID2 immunofluorescence (Figure 4B). Moreover, streptavidin labeling confirmed presence of biotinylated signal overlapping with BioID2 immunoreactivity confirming occurrence of local biotinylation. Next AAV2/9 viruses containing this construct were injected in the cerebral cortices of neonatal mice, and after 5 weeks, the substrate, biotin, was administered through intraperitoneal injection for seven consecutive days (Figure 4A). At 6 weeks of age, mice where sacrificed and biotinylated proteins were purified using streptavidin beads and identified by mass spectrometry. As a control, in parallel experiments, a fusion protein of BioID2 and Shank3* (a fragment of Shank3 comprising amino acids 1055-1806 to make it packageable in AAV) (Figure 4A) was expressed in mouse cerebral cortices by the same procedure. Like synaptopodin, Shank3* is an actin associated protein that is enriched in dendritic spines, but concentrated in the subsynaptic region, rather than close to the spine neck where the synaptopodin is localized. Accordingly, streptavidin labeling of cells expressing BioID2-Shank3* revealed that the biotin signal generated by this construct was juxtaposed, rather than overlapping, with mRFP-synaptopodin. Thus, comparison of the proteome of BioID2-synaptopodin and BioID2-Shank3* is expected to allow for spatial mapping of the dendritic spine and to identify proteins specifically localized close to synaptopodin (Figure 4C). We performed two separate experiments in which brain extract of multiple pups were collected and results were statistically analyzed to determine the proteins that are significantly enriched in SA-targeted samples (see methods for details).

**Figure 4.**
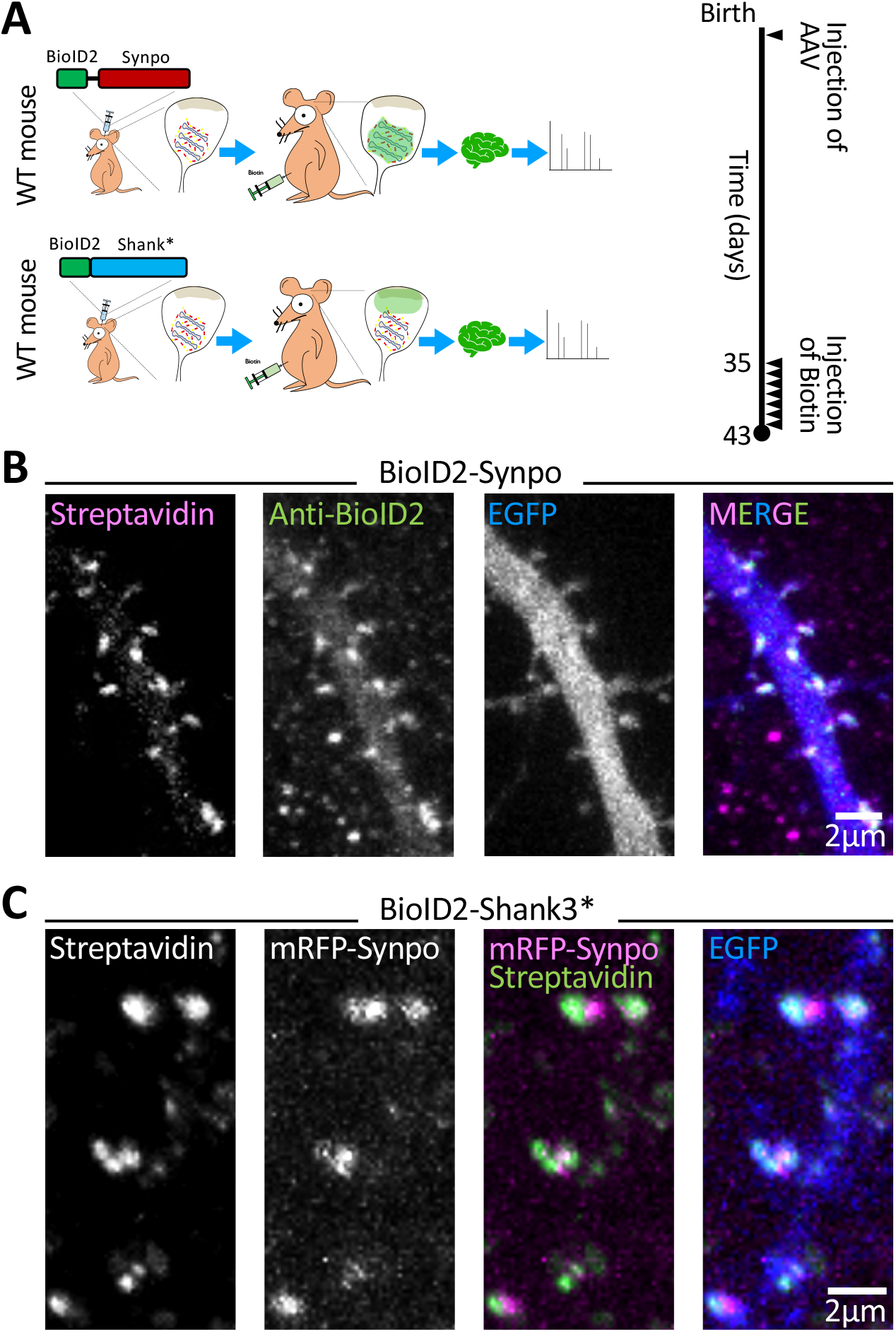
Identification of SA associated proteins by in vivo proximity labeling. **A**. Schematic representation of in vivo proximity labeling of protein neighbors of synaptopodin (iBioID). To target the biotinylating enzyme to the spine apparatus, AAV2/9 viruses containing BioID2-synaptopodin were injected in the cortex of neonatal mice. BioID2 fused to a fragment of Shank3 (BioID-Shank3*) was used as a control to localize the biotinylating enzyme to a different region of the spine, i.e., the neighborhood of the post-synaptic density. Biotin was administered IP starting five weeks after birth, for seven consecutive days. On day 8 the mouse was sacrificed, and biotinylated proteins were isolated from their brain. **B**. Cultured hippocampal neurons transfected with BioID2-synaptopodin and EGFP were stained with Alexa647-Streptavidin and anti-BioID2 antibody to confirm the specificity of biotinylation in spines. **C**. A cultured hippocampal neuron expressing mRFP-synaptopodin, BioID2-Shank3* and EGFP were labeled with Streptavidin to examine the specificity of biotinylation, as well as the differential localization of synaptopodin and of the control protein Shank3* in spines.

We identified 140 proteins enriched with statistical significance in the bioID2-synaptopodin proteome relative to the Shank3* proteome. These included a large number of signaling and scaffold proteins as well as actin-related likely comprising structural components of the SA as well as “client” and regulatory proteins. Consistent with an association of synaptopodin with the ER, they also included many ER components, such as proteins involved in calcium storage and signaling (e.g., Ryr2, Ip3r3, and Stim1), lipid transfer (e.g., Pitpnm1-3, Osbpl and TMEM24/C2cd2l) (Figure S3) and lipid metabolism (for example Faah, a protein implicated in endocannabinoid metabolism and previously reported to be localized in spines (Borgmeyer et al., 2021)). As expected, the BioID2-Shank3* control proteome was more enriched in proteins of post-synaptic densities, such as Shank1-2 and Homer1-3 and receptors for glutamate, including the NMDA receptors Grin1, Grin2a and Grin2b, the AMPA receptors Gria2 and Gria3 as well as the metabotropic receptors Grm2 and Grm5 (Table S1). The difference between the two proteomes was further supported by the analysis of Gene Ontology terms for the identified proteins. The highest enriched cellular components for BioID2-synaptopodin samples included ER tubular network and cortical ER, while the enriched GO-terms for BioID2-Shank3* samples included anchored and cytosolic components of post-synaptic densities. Even the actin associated proteins enriched in the two proteomes were different, as BioID2-synaptopodin samples were enriched for contractile actin and stress fiber components (e.g., α-actinin-1, α-actinin-2, and α-actinin-4, Myosin-9), while BioID2-Shank3* samples contained components of Arp2/3 and SCAR/WAVE complexes (e.g., Arpc1a, Arpc2, Arpc5l, Wasf3, Abi1-2, Nckap1, and Cyfip1-2).

### Colocalization of identified proteins with synaptopodin in dendritic spines

To validate our biochemical findings, we investigated whether a selected few identified proteins are indeed localized in close proximity of synaptopodin in situ. We focused on proteins with a proposed or proven link to synaptopodin such as Pdlim7 (a member of the PDZ and LIM domain protein family and a potential interactor reported in <u>Biogrid</u>, (https://thebiogrid.org/), Magi1 (a member of the MAGUK protein family that comprises WW, PDZ and GK domains (Patrie et al., 2002), α-actinin-2 and α-actinin-4 (Asanuma et al., 2005; Kremerskothen et al., 2005) and Magi2 (a putative synaptopodin interactor because of its similarity to Magi1) (Nagashima et al., 2015a). To this aim, we expressed these proteins with a fluorescent tag in cultured hippocampal neurons. We found that in dendritic spines and at axonal initial segments, EGFP-Pdlim7 precisely colocalized with mRFP-synaptopodin (Figures 5A and S4A). Consistent with previous reports that α-actinin-2 colocalize with synaptopodin at the same sites, strong overlap was also observed between synaptopodin and EGFP-α-actinin-2 in spines. EGFP-Magi1 and EGFP-Magi2 (S-SCAM) also colocalized with mRFP-synaptopodin in dendritic spines (Figure 5B-C). However, these two proteins were not enriched at axonal initial segment (Figure S4B-C). The different appearance of spines upon exogenous expression of Magi1 and Magi2 proteins (fewer and larger) and of α-actinin (thinner and longer) is in agreement with previous reports (Danielson et al., 2012; Nakagawa et al., 2004). However, synaptopodin was not required for the localization of EGFP-Magi1/2 and EGFP-α-actinin-2 in dendritic spines, as these proteins still localized in spines in synaptopodin KO hippocampal cultured neurons (Figure 5F-H). In contrast, in synaptopodin KO neurons, an obvious greater cytosolic pool of Pdlim7 throughout dendrites was observed (Figure 5E), pointing to a role of synaptopodin in enriching Pdlim7 in spines and thus to a special relation between synaptopodin and Pdlim7.

**Figure 5.**
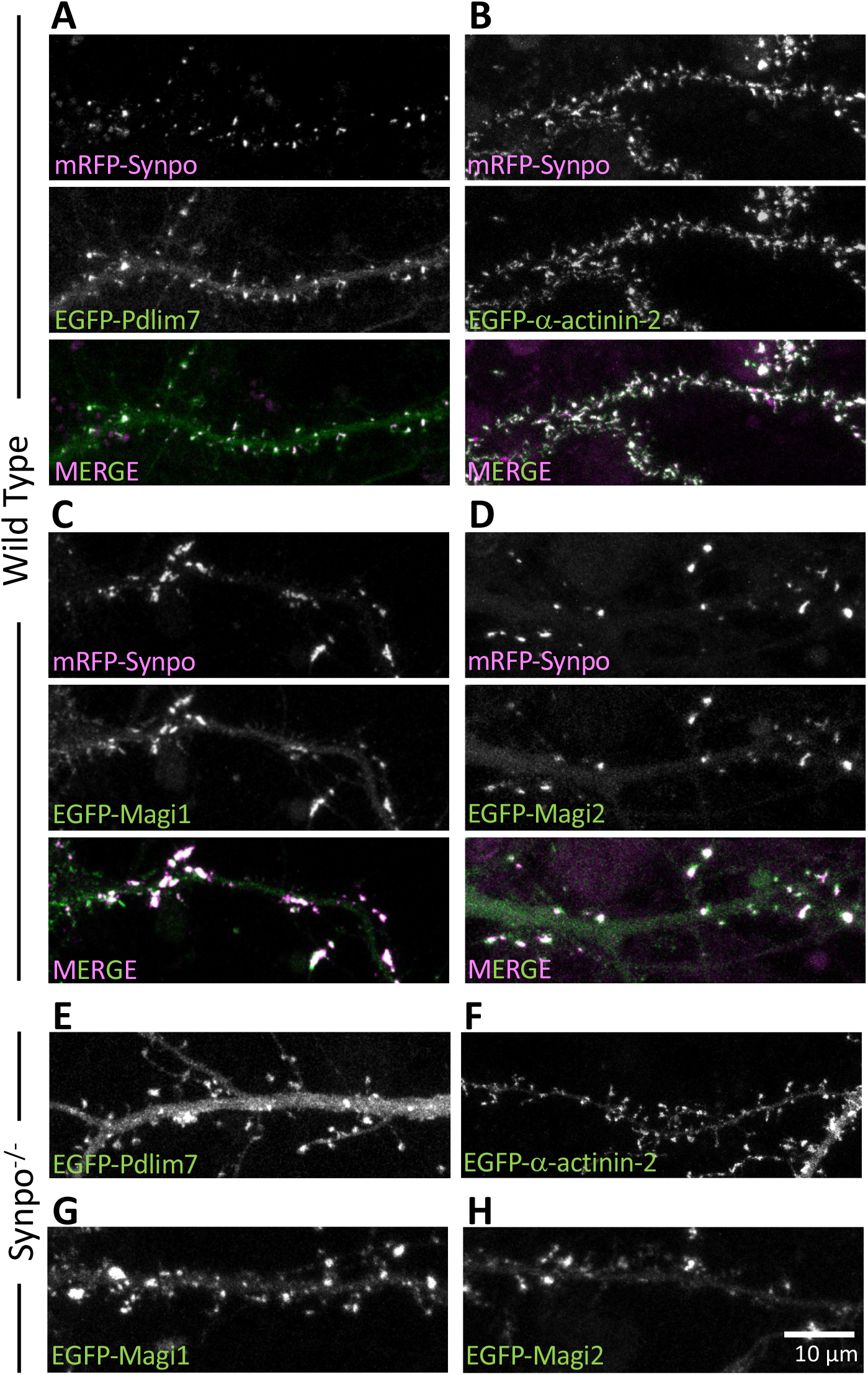
Co-localization with synaptopodin in dendritic spines of proteins identified by our screen. **A-D**. Cultured WT hippocampal neurons expressing mRFP-synaptopodin with either EGFP-Pdlim7 (**A**), EGFP-α-actinin2 (**B**), EGFP-Magi1 (**C**) or EGFP-Magi2 (**D**). **E-H**. Cultured hippocampal neurons of synaptopodin KO mice transfected with EGFP-Pdlim7 (**E**), EGFP-α-actinin-2 (**F**), EGFP-Magi1 (**G**), or EGFP-Magi2 (**H**).

### Interaction of synaptopodin with Magi1, Magi2 and Pdlim7 as assessed by expression in non-neuronal cells

We further investigated the relation of Pdlim7, Magi1 and Magi2 to synaptopodin by expressing them in non-neuronal cells. When expressed alone in COS-7 cells, either EGFP-Magi1 or EGFP-Magi2 localized under the plasma membrane with an enrichment at cell-cell junctions (Figures 6A and 6C), but upon coexpression with synaptopodin these two proteins were also recruited to synaptopodin positive internal structures (Figures 6B and 6D). Similarly, expression of the ER-targeted chimera (synaptopodin-ER) resulted in the relocation of Magi1 and Magi2 to synaptopodin assemblies at ER (Figure S4E-F).

**Figure 6.**
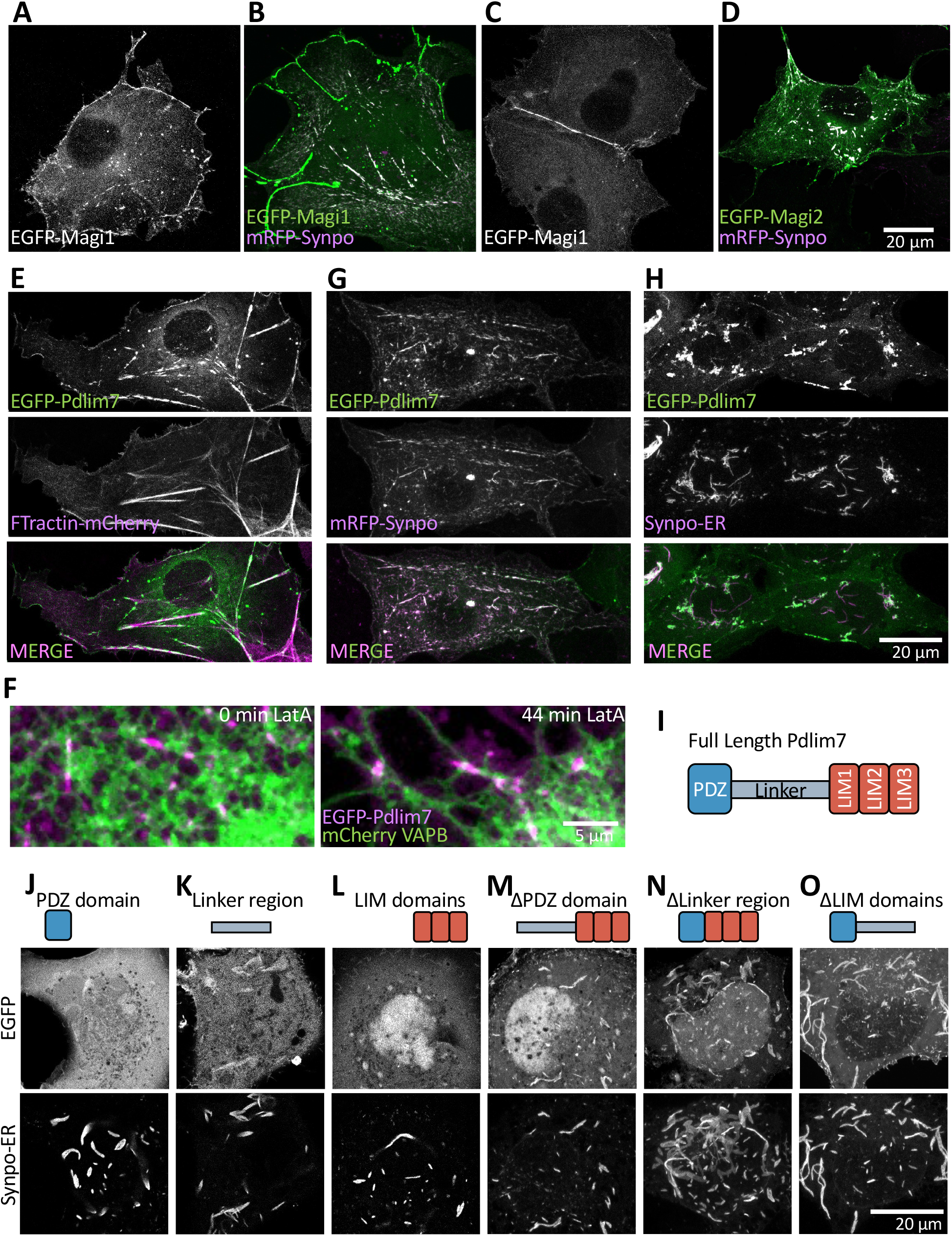
Proteins identified by our screen interact with synaptopodin in COS-7 cells. **A-D**. EGFP-Magi1 and EGFP-Magi2 localize under the cell surface and primarily at cell-cell junctions when expressed alone in COS-7 cells. Both proteins partially relocalize to synaptopodin-positive structures when expressed together with mRFP-synaptopodin. **E**. EGFP-Pdlim7 partially colocalizes with FTractin-mCherry on stress fibers. **F**. A representative COS-7 cell expressing EGFP-Pdlim7 (magenta) and mCherry-VAPB (green) before and after treatment with 2µM LatA, showing that EGFP-Pdlim7 on stress fibers collapses onto the ER after treatment with LatA. **G**. mRFP-synaptopodin precisely colocalizes with EGFP-Pdlim7. **H**. Coexpression of synaptopodin-ER with EGFP-Pdlim7 results in recruitment of Pdlim7 to synaptopodin assemblies in the ER. **I**. Domain structure of Pdlim7. **J-O**. Co-expression of the Pdlim7 fragments shown with synaptopodin-ER. All Pdlim7 constructs were tagged with EGFP at N-term. All constructs containing the linker region colocalize with synaptopodin-ER. While the PDZ domain or the LIM domains alone show little colocalization with synaptopodin-ER, a construct comprising both regions interacts with synaptopodin even in the absence of the linker region. This data demonstrates the multivalency in the interaction between synaptopodin and Pdlim7.

In agreement with its reported role in the regulation of the actin cytoskeleton (Krcmery et al., 2013; Urban et al., 2016), EGFP-Pdlim7 was associated with actin stress fibers when expressed alone in COS-7 cells, as indicated by its colocalization with FTractin-mCherry (Figure 6E), and showed a discontinuous localization on them, as observed for synaptopodin. Moreover, similar to synaptopodin, EGFP-Pdlim7 associated with stress fibers collapsed onto the ER upon LatA treatment (Figure 6F, Video 4). When co-expressed with mRFP-synaptopodin, EGFP-Pdlim7 precisely colocalized on all structures positive for this protein, including ones insensitive to 2µM LatA (Figures 6G and S5, Video 5). It also precisely colocalized with the ectopically targeted synaptopodin (synaptopodin-ER) (Figure 6H). As Pdlim7 is a newly identified component of spines and is the protein that precisely colocalize with synaptopodin in both neurons and fibroblasts, we examined in more detail its interaction with synaptopodin. Coexpression of Pdlim7-deletion constructs with synaptopodin revealed that all constructs containing the linker region, including the linker region alone, robustly colocalized with synaptopodin-ER (Figure 6K, 6M and 6O). However, while the LIM domain region alone (Figure 6L) and the PDZ domain alone (Figure 6J) were primarily cytosolic or nuclear, respectively, a construct comprising the LIM domains and the PDZ domain, but lacking the linker region, also colocalized strongly with synaptopodin (Figure 6N). We conclude that interactions involving multiple domains of Pdlim7, i.e., a strong interaction with the linker region and weak interactions that function cooperatively within the PDZ and the LIM domains, drive its co-assembly with synaptopodin.

Finally, when synaptopodin, Magi1, Magi2 and Pdlim7 were all coexpressed together in COS-7 cells, they all co-assembled into the same structures (Figure 7A). Interestingly, synaptopodin was not necessary for this coassembly, as BioID2-Pdlim7 was sufficient to recruit Magi1 and Magi2 (Figure 7B). These results underscore the multivalency of the interactions among identified SA-associated proteins, consistent with a model in which phase separation helps drive the assembly of the cytosolic components of SA.

**Figure 7.**
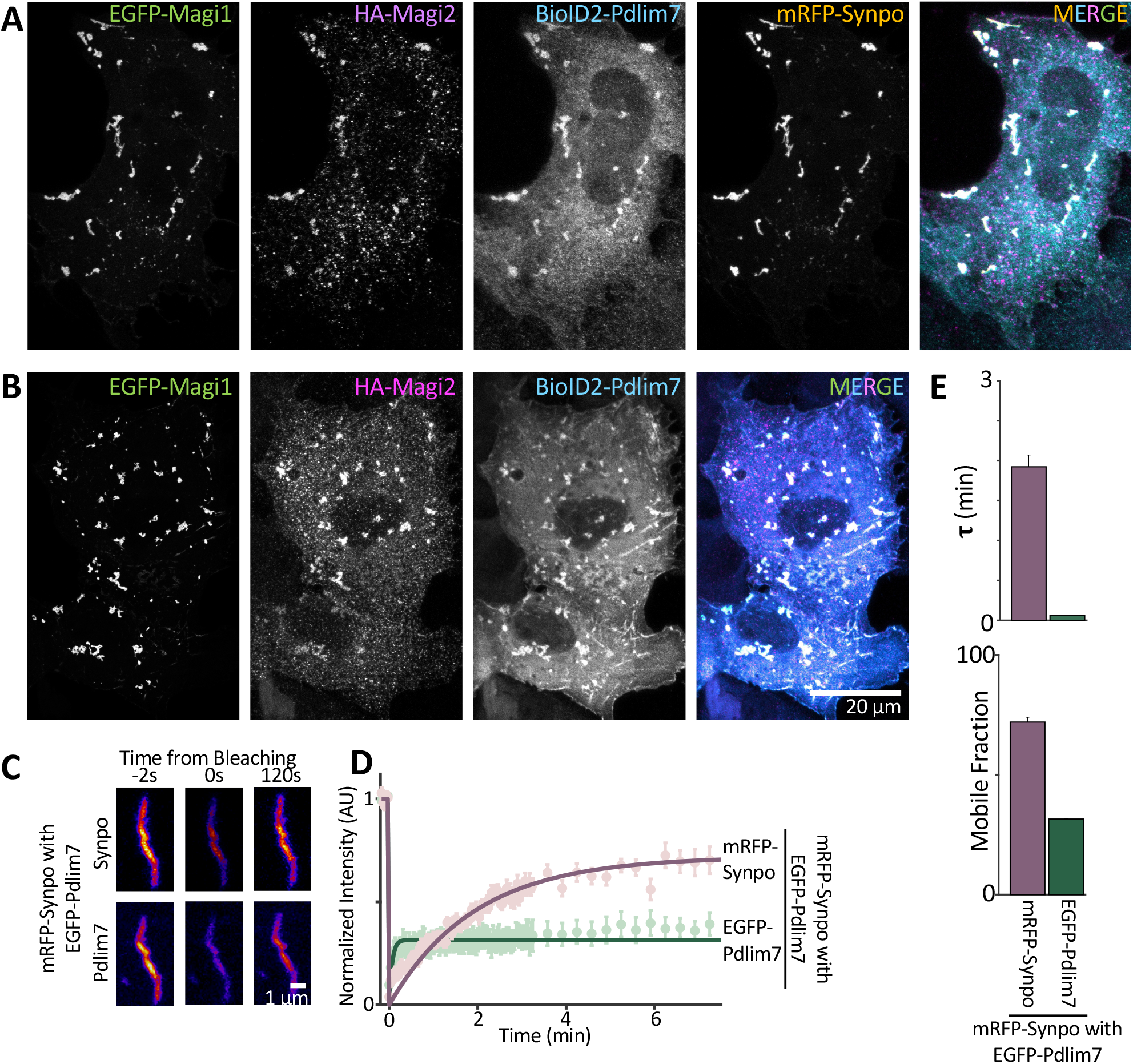
Pdlim7 stabilizes synaptopodin assemblies. **A**. When mRFP-synaptopodin, EGFP-Magi1, HA-Magi2 (labeled with anti-HA antibody) and BioID2-Pdlim7 (labeled with anti-BioID2 antibody) are coexpressed in COS-7 cells, they all colocalize. **B**. Even in the absence of synaptopodin, Pdlim7 can recruit Magi1 and Magi2 to internal structures. **C-E** The dynamics of synaptopodin, as examined by FRAP in COS-7 cells, changes in the presence of Pdlim7. **C** Representative images of the assemblies containing both mRFP-synaptopodin and EGFP-Pdlim7 before bleaching (time -2s) and at 0 and 2 min after photobleaching in COS-7 is shown. Quantifications of the recovery are shown in **D** and **E** (n = 15). The lines in D show exponential fits to the fluorescent recovery curve. All intensities are normalized to the average intensity in the first five images pre-bleaching. See methods for more details. **E**. The recovery time constant and the mobile fraction was calculated for each condition from the fitted single exponential curves.

### Pdlim7 stabilizes synaptopodin assemblies

To further investigate the impact of Pdlim7 on synaptopodin assemblies, we examined the impact of its presence on synaptopodin condensates. To this aim we focused on non-tress fiber assemblies. Coexpression with Pdlim7 strongly slowed down the recovery of synaptopodin, as only 46% of its fluorescent signal recovered within the first two minutes after photobleaching (Figure 7 C-D), compared to 77% when synaptopodin was expressed alone (Figure 3D). The mobile fraction of synaptopodin was also reduced from 91.3 ± 0.6% to 72 ± 2% (Figure 7E and 3F). The time constant, τ, of recovery increased from 1.05 ± 0.03 to 1.9 ± 0.1 min (n = 15) (Figure 7E and 3E). When EGFP-Pdlim7 in the same assemblies was photobleached, 69.5 ± 0.1% of the fluorescent signal did not recover within the seven minutes of the experiment, indicating that the bulk of EGFP-Pdlim7 in these assemblies are immobile within the time frame of our experiment (Figure 7D). The remaining 31% recovers very fast with a τ of 3.6 ± 0.24 s, suggesting that they are in the unbound and/or cytosolic state. These results indicate that Pdlim7 may function as a molecular anchor to stabilize the assembly of SA proteins.

### Analysis of proteins with expression pattern similar to synaptopodin

Proteins that are part of the same structure or cooperate in their function may have similar patterns of expression. The expression pattern of synaptopodin varies depending on the neuronal population. Its highest level of expression is in cortical and hippocampal neurons where the SA is also detectable, while neurons such as Purkinje cells that lack the SA, despite having a high abundance of dendritic spines, do not express synaptopodin. So, as a complementary approach to proximity biotinylation, we analyzed available single-cell RNA-seq datasets to find proteins with an expression pattern similar to that of synaptopodin. We used the number of reads for different genes from mouse brain cells available on DropViz (Saunders et al., 2018) and compared the expression pattern of any single gene of interest (GoI) with that of synaptopodin in all cells of the dataset. To quantify the correlation between the pattern of expression of synaptopodin with any other GoI, we calculated the Pearson correlation coefficient, *r*, using the following equation:

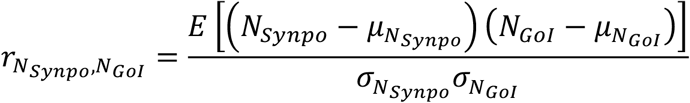

where *N*_*Synpo*_ and *N*_*GoI*_ are the number of reads of synaptopodin and GoI in each cell, *E* is the expected value, 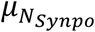 and 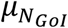, are the means of reads in each cell for synaptopodin and for every gene, respectively, and 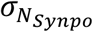 and 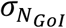 are the standard deviations of *N*_*Synpo*_ and *N*_*GoI*_, respectively. These correlation values vary between -0.037 and 0.558 (Figure S6A). The 125 genes (among the total 32307 genes of the database) whose expression patter correlated with that of synaptopodin with an r value greater than 0.2 are shown in Figure S6B. Bioinformatic analysis of the identified proteins shows that the most enriched GO-terms are postsynaptic cytoskeleton organization, regulation of ion transport, and regulation of chemical synaptic transmission, consistent with the role of the SA at post-synapse. Interestingly, Pdlim7, identified above as a hit by proximity biotinylation, was part of this list, confirming that this protein is a major functional partner of synaptopodin.

## DISCUSSION

While the spine apparatus has been observed for over 70 years, the mechanism of its formation and function remains elusive mainly due to the lack of a comprehensive understanding of its molecular composition. More specifically, the nature of the dense matrix connecting ER cisterns remains unknown, besides evidence that it contains the protein synaptopodin, an actin binding protein. The findings reported here suggest that synaptopodin interacts (directly or indirectly) with the ER membrane, although the underlying mechanism(s) remain unclear. Importantly, we identify 140 proteins with a propensity to localize in proximity of synaptopodin in brain, as assessed by their enrichment in the proximity proteome of BioID2-synaptopodin relative to the proximity proteome of BioID2-Shank3*, a molecular component of the post-synaptic density. We confirmed the colocalization and selective concentration of four of the hits identified by this analysis, Pdlim7, Magi1, Magi2 and α-actinin-2 with synaptopodin in neuronal dendritic spines (Asanuma et al., 2005; Kremerskothen et al., 2005; Nagashima et al., 2015a; Patrie et al., 2002; Sánchez-Ponce et al., 2012). Furthermore, we showed that when Pdlim7, Magi1, and Magi2 are expressed together in non-neuronal cells they co-assemble with synaptopodin and actin, supporting the possibility that they may be part of the matrix that connects to each other ER cisterns of the SA. We additionally provide evidence for a liquid nature of the protein assemblies involving synaptopodin, suggesting that the matrix connecting ER cisterns may self-organize according to phase separation principles. In a complementary approach, we searched for proteins whose expression pattern in brain is similar to the expression pattern of synaptopodin. The list of approximately 120 proteins identified by this search included Pdlim7, thus supporting the likely role of Pdlim7 as a critical functional partner of synaptopodin.

The different proximity proteome of BioID2-synaptopodin and BioID2-Shank3* validates the in vivo approach that we have employed toward the identification of components of the SA and of proteins that may be concentrated in its proximity within spines. As expected, these two proteomes comprise transmembrane proteins, but primarily ER proteins in the case of synaptopodin, and plasma membrane proteins in the case of Shank3*. Analysis of the synaptopodin proteome reveals a variety of house-keeping proteins of the smooth ER, but none of the most enriched ER proteins that we tested were found to be specifically concentrated in dendritic spines. Thus, we have found no evidence that the stacking of ER cisterns in the SA reflects the concentration at these sites of a specific ER protein that interacts with synaptopodin or with another protein of the intervening matrix, although the existence of such protein remains possible. A plausible scenario is that the stacking of ER-cisterns via an intervening protein matrix may be due to a multiplicity of low affinity interactions between components of this matrix and the ER membrane.

The protein neighbors of synaptopodin that we have validated are all factors implicated in actin-based scaffolds. Pdlim7, an actin regulatory protein known to have an important role in a variety of cell types (D’Cruz et al., 2016; Krcmery et al., 2013; Urban et al., 2016), was a previously unknown component of the SA. Its striking colocalization with synaptopodin both in neurons and in fibroblasts, as well as the similar pattern of expression of the two proteins in brain, suggests their critical functional partnership. Presence of the PDZ domain in Pdlim7 would be consistent with an interaction with a membrane protein (Ye and Zhang, 2013), but so far binding partners for this PDZ domain have not been identified. Most frequently, PDZ domains bind to plasma membrane localized proteins(Ye and Zhang, 2013), but it cannot be excluded that Pdlim7 may anchor the matrix to some intrinsic membrane protein of ER cisterns.

MAGUK inverted (Magi) proteins Magi1 and Magi2 (S-SCAM) are scaffolding proteins typically localized in the protein matrix that lines the cytosolic leaflet of the plasma membrane, primarily at cell-cell junctions. In brain, Magi2 was shown to be a structural component of post-synaptic densities that regulate the trafficking of NMDA and AMPA receptors (Danielson et al., 2012; Deng et al., 2006; Emtage et al., 2009; Hirao et al., 1998; Nagashima et al., 2015a). Thus, the enrichment of Magi2 (S-SCAM) in the BioID2-synaptopodin proteome, relative to BioID2-Shank3* proteome, was surprising. However, its paralog Magi1 is a known interactor of synaptopodin in kidney podocytes (Patrie et al., 2002) and both Magi1 and Magi2 were shown to play a role in the organization of the glomerular filtration barrier, consistent with the known role of synaptopodin in the generation and maintenance of such barrier (Nagashima et al., 2015; PATRIE et al., 2001; Yamada et al., 2021). The interaction of Magi1 and Magi2 with synaptopodin is supported by our transfection experiments in non-neuronal cells, where both Magi1 and Magi2 localized under the plasma membrane and predominantly at cell-cell contacts when expressed alone, but partially relocalized to internal synaptopodin positive structures when co-expressed with synaptopodin or Pdlim7. The matrix of the SA may contain a reserve pool of these proteins. In fact, acting as a reservoir of proteins implicated in post-synaptic function may be one of the functions of the SA.

The idea that principles of phase separation may apply to the formation of the matrix connecting cisterns of the SA is in line with what is reported for other membrane associated scaffolds, including focal adhesions (Ramella et al., 2021; Wang et al., 2021), pre- and postsynaptic densities (Araki et al., 2016; Milovanovic et al., 2018; Park et al., 2021; Wu et al., 2021) and receptor-associated signaling scaffolds (Su et al., 2016). The liquid organization of a synaptopodin-based matrix is supported 1) by evidence that the fluorescence of synaptopodin assemblies can quickly recover after photobleaching and 2) by the observation that such assemblies expand in a reversible fashion by lowering the temperature, revealing a thermodynamically-driven process. Macromolecular condensates generated by phase separation rely on protein self-assembly by intrinsically disordered regions and/or on the multivalency of the interactions of participating proteins. Synaptopodin is predicted to be a primarily disordered protein and its interactors discussed here are multimodular proteins. Moreover, as we have shown for Pdlim7, its binding to such protein involves multiples domains of Pdlim7.

Previous reports showed that the presence of synaptopodin assemblies is necessary for the stability and long-term survival of dendritic spines (Okubo-Suzuki et al., 2008; Yap et al., 2020b). Similarly, the ER is more consistently present in the spines that contain synaptopodin (Perez-Alvarez et al., 2020). Presence of Pdlim7 in mRFP-synaptopodin assemblies delays the recovery of mRFP fluorescence after photobleaching, revealing that it increases the stability of such assemblies. Other SA proteins may have additional stabilizing effects, thus contributing to the stability of the SA.

Although the focus of the present study was to elucidate mechanisms underlying the structure of the spine apparatus, the synaptopodin and Shank3* proximity proteome that we have characterized can be mined to learn more about spine cell biology. Our results also showed that the in vivo proximity labeling approach employed by us, and originally used to identify proteins implicated in post-synaptic inhibitory signaling (Uezu et al., 2016b), is a very powerful methodology to study subdomains of the dendritic spine. In addition, the correlative analysis of single-cell RNA-seq data provided insights for future studies of regulatory mechanisms relevant to SA biology. Taken together, the data provided here is a foundation not only for understanding the formation and function of the SA, but also for gaining new mechanistic insight into physiological processes occurring in dendritic spines.

## ACKNOWLEDGEMENTS

We thank members of the De Camilli lab for advice and discussion. This work was supported in part by NIH grant NS36251, DK45735, the HHMI and the Kavli Foundation to P.D.C. H.F. was supported by an HHMI/Life Sciences Fellowship.

## AUTHOR CONTRIBUTIONS

P.D.C. and H.F. designed experiments. Y.W. performed EM experiments. H.F. and V.F. performed characterizations of Pdlim7. All other experiments and analyses were performed by H.F. All authors interpreted the data. H.F. and P.D.C. wrote the manuscript with input from all authors.

## MATERIALS AND METHODS

### Antibodies and Reagents

The list of antibodies, their working dilution, and the supplier for this study can be found in **Table S2**. pAAV-HA-GFP-Pdlim7, pAAV-HA-Magi1 and pAAV-HA-Magi2 were cloned by Genscript. AAV2/9 packaging for pAAV-BioID2-synaptopodin was done by Janelia Virus Services, HHMI, and packaging of pAAV-BioID2-Shank3^*^ was done by Penn Vector Core, University of Pennsylvania. The following constructs were kind gifts: MCS-BioID2-HA from K. Roux (Addgene plasmid #74224); pCDNA flag MAGI1c from W. Sellers (Addgene plasmid #10714); Myc rat S-SCM from Y. Hata and Y. Takei (Addgene plasmid #40213); EGFP-α-actinin-2 from J. Hell (Addgene plasmid #52669); EGFP-MyosinIIA from M. Krummel (Addgene plasmid #38297); F-Tractin-mCherry from T. Meyer(Addgene #155218); pAcGFP-Sec61ß from E. Schirmer (Addgene plasmid #62008) ; mRFP-synaptopodin from A. Triller (Institut de Biologie de l’Ecole Normale Supérieure (IBENS), Paris, France); RFP-Shank3 from A. Koleske (Yale School of Medicine, New Haven, CT, USA). pAAV-GFP-MCS, Halo-Sec61ß, EGFP-MOSPD1, and EGFP-VAPB were previously constructed in our lab.

Biotin was purchased from Sigma (B4501). Latrunculin A and Rock inhibitor Y-27632 were from EMD Millipore Corp.

### Generation of Plasmids

Most constructs were generated with regular cloning protocols or through site-directed mutagenesis. Some constructs were ligated using Gibson assembly (NEB) or In-Fusion Cloning (Takara Bio). Details of primer sets, enzymes, techniques, and plasmids used for each construct can be found in **Table S3**.

The desired ORF for Pdlim7 (Gene bank ID: AF345904.1) was amplified by PCR from Human Universal QUICK-Clone™ II (Takara 637260) using primers depicted in Table S2 and inserted into pEGFP-C1 plasmid through Gibson assembly.

All constructs were sequenced in their entirety before use in any experiment.

### Semi-Automated detection of SA in FIB-SEM images

To morphologically characterize SA in FIB-SEM images, an in-house ImageJ macro was developed and used. First a Difference of Gaussian (DoG) filter was applied to the images to detect the edges of features of interest. To generate a mask, a manually selected threshold was applied to the filtered image. Then the mask for the feature of interest was selected in one plane manually and tracked automatically in other planes. This automatic tracking takes advantage of the continuity in the structure of the SA. The final mask can also be refined by adding or removing mischaracterized portions of the mask, using a similar semiautomated approach. Other subcellular features such as plasma membrane, PSD or mitochondria can be detected with this code by changing the DoG filter.

### In Vivo BioID (iBioID)

iBioID was performed using a method described previously (cite) with some minor modifications. The AAV2/9 viruses containing pAAV-BioID2-synaptopodin or pAAV-BioID2-Shank3* were injected into C57/B6J mouse cortex at P0 to P2. At five weeks of age, biotin was subcutaneously injected at 24 mg/kg for 7 consecutive days to increase the biotinylation efficiency. Biotinylated proteins were purified from the forebrain of each mouse separately. This experiment was performed on two different litters. To better control for the genetic variability, each round of experiment included injections of all the pups of the litter. In the first case (experiment 1), 4 mice were injected with BioID2-synaptopodin and 3 with BioID2-Shank3*. In the second experiment (experiment 2), 5 mice were injected with BioID2-synaptopodin and 4 with BioID2-Shank3*. Biotinylated proteins from each litter were purified using two slightly different methods.

For experiment 1, the forebrains were homogenized on ice using glass/Teflon homogenizer with 0.5ml of lysis buffer (50mM HEPES pH 7.5m 150mM of NaCl, 1mM EDTA, 0.25µl benzonase, EDTA-free cOmplete mini protease inhibitor cocktail -Roche Diagnostics 11836170001). The samples were then transferred to an Eppendorf tube, and lysis buffer containing 10% SDS was added to reach a final SDS concentration of 1% and incubated at 50°C for 15 min on a shaking thermal block. Next Lysis buffer containing 20% Triton-X100 was added to reach a final concentration of 2% for Triton-X100. The samples were then incubated on ice for 5 minutes. 10µls of each of the samples were separated for analysis with Western blot. The remaining portions of each sample were incubated overnight at 4°C with 250µg of Pierce Streptavidin magnetic beads (S-beads) that were pre-equilibrated with lysis buffer.

For experiment 2, the forebrains were also homogenized on ice using glass/Teflon homogenizer with 3ml of lysis buffer (50mM HEPES pH 7.5m 150mM of NaCl, 1mM EDTA, 0.25µl benzonase, EDTA-free cOmplete mini protease inhibitor cocktail -Roche Diagnostics 11836170001). The samples were then transferred to an Eppendorf tube, and lysis buffer with 10% SDS was added to reach a final SDS concentration of 1% and incubated at 50°C for 15 min on a shaking thermal block. Triton-X100 was not added in this experiment. 10µls of each of the samples were separated for analysis with Western blot. The remaining portions of each sample were incubated for 2hrs at room temperature with 250µg of Pierce Streptavidin magnetic beads (S-beads) that were pre-equilibrated with the lysis buffer.

In both experiments, the S-beads incubated with samples where then collected with a magnetic stand and the supernatant was discarded. S-beads were washed as follows: 2x with 300 µl of wash buffer 1 (50mM HEPES pH 7.5, 150mM NaCl, 1mM EDTA, 2%SDS), 2x with wash buffer 2 (50mM HEPES pH 7.5, 1% Triton X-100, 1% deoxycholate, 25mM LiCl), 2x with wash buffer 3 (50mM HEPES pH 7.5, 1M NaCl), 5x with wash buffer 4 (50mM ammonium bicarbonate). The bound proteins were then eluted by incubating the beads for 1.5h at 60°C with 100 ul of elution buffer (50mM ammonium bicarbonate, 0.1% Rapigest SF surfactant, 2mM biotin), followed by a second round or elusion with 100µl of elution buffer for 1.5h at 60°C. The final protein concentration was measured by BCA. Liquid Chromatography and Mass Spectrometry was performed by W.M. Keck Foundation Biotechnology Resource Laboratory (Yale School of Medicine, New Haven, CT).

### Statistical Analysis of the identified biotinylated proteins

To identify the proteins that were present at significantly different concentration in the BioID2-synaptopodin samples relative to the BioID2-Shank3* samples, we performed statistical analysis. The material from each mouse brain was considered one biological replicate. The goal of this analysis was to correct for the variability between different samples, in terms of expression level of the BioD2-fusion protein and of the efficacy of the fusion protein in performing biotinylation. In experiment 1, we measured the enrichment of any given protein in each group as the ratio between mean number of peptides in the BioID2-synaptopodin samples relative to BioID2-Shank3* samples. To determine the significance of enrichment, we performed a student’s t-test between the samples of the two groups and defined a p-value < 0.1 as significant enrichment.

In experiment 2 we performed an additional normalization to correct for the variation in the yield of biotinylation between BioID2-synaptopodin and BioID2-Shank3*, and among different animals. To this aim, we used the two major endogenously biotinylated proteins, Propionyl-CoA carboxylase alpha chain (Pcca) and pyruvate carboxylase (Pc), as an internal control. Higher number of reads for these endogenous proteins relative to proteins biotinylated by exogenous BioID2 meant lower biotinylation yield. Subsequently, we normalized the amount of the identified proteins for each sample to the sum of the number of reads for these two proteins, and the normalized values for synaptopodin and Shank3* were compared as described above for the first experiment.

### Determining the correlative expression of proteins relative to synaptopodin

To find the genes with an expression pattern similar to that of **synaptopodin**, the number of reads per protein per mouse brain cell was obtained from Dropviz (cite). An inhouse R code was developed to calculate the Pearson correlation coefficient for individual genes with synaptopodin according to equation 1 (see main text). Due to the large sample size (n = 939489 cells) and according to equation S1, we estimate that non-zero correlation coefficients can be considered any value above 10^−3^, as they will have a t of 0.99 in the Student’s t-distribution.

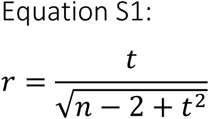

where n is the number of samples, 2 is the degrees of freedom assuming a normal sample distribution and r is the Pearson correlation coefficient. We set the artificial threshold of r = 0.2 to determine the proteins with expression pattern similar to synaptopodin.

### Primary Neuronal culture and transfection

Hippocampus of P0-P1 C57BL/6 mice (Jackson Laboratory) were dissected on ice in Hibernate-A media (Thermofisher). Tissues were then washed in ice cold Dissociation Media (5.8mM MgCl_2_, 0.252mM CaCl_2_, 10mM HEPES pH = 7.4, 1mM Pyruvic Acid, 81.7% Mg and Ca free HBSS from Thermofisher, in water) and immediately digested in Cystein-activated Papain solution (17U/ml Papain from Worthington, 20µg/ml DNase I from Sigma, 2mg/ml of L-Cysteine Hydrochloride from Sigma in Dissociation media) for 30 min at 37°C. Papain was then inactivated with 10% FBS (in Dissociation media), followed by washes in Dissociation media and in Neurobasal-A (Thermofisher) supplemented with 2% B27 and 2mM L-Glutamax. 120-150K cells were plated on the glass bottom of Mattek plates in 150µl of Neuronal Growth Media (Neurobasal-A supplemented with 2% B27, 2mM L-Glutamax, 15% gilial enriched media and 10% cortical enriched media). 4-16hrs after plating, 2ml of Neuronal Growth Media was added per plate. 0.5ml of Neuronal Growth Media was added per dish every 3-4 days afterwards. Hippocampal neurons were transfected on DIV11-13 using CalPhos Mammalian Transfection kit (Takara) per manufacturer’s instructions, and fixed or imaged at DIV16-24.

### Non-neuronal Cell Cultures and Transfections

hTERT-RPE1 cells were a kind gift of A. Audhya (University of Wisconsin, Madison, WI). COS-7 cells were obtained from ATCC. Cells were maintained at 37°C in humidified atmosphere at 5% CO_2_. COS-7 cells were grown in DMEM and RPE1 cells in DMEM/F12 medium (Thermo Fisher Scientific) supplemented with 10% FBS, 100 U/mL penicillin, 100mg/mL streptomycin, and 2mM glutamax (Thermo Fisher Scientific). All cell lines were routinely tested and always resulted free from mycoplasma contamination.

The cells were seeded on glass bottom mat-tek dishes at least 16 hours before transfection. All transfections of plasmids used Lipofectamin 2000 (Thermofisher) to manufacturers specifications for 16-24 hours in complete media.

### Live Cell Imaging and Immunofluorescence

Confocal imaging was performed using LSM880 or LSM800 (Carl Zeiss Microscopy) with a 63X/1.40 NA plan-apochromat differential interference contrast (DIC) oil immersion objective and 32-channel gallium arsenide phosphide (GaAsP)-photomultiplier tubes (PMT) area detector. 405nm, 488 nm, 561 nm and 633 laser lines were used in this study.

For live imaging, non-neuronal cells were imaged using Live Cell Imaging buffer (Life Technologies), and cultured neurons were imaged in modified Tyrode Buffer (119mM NaCl, 5mM KCl, 2mM CaCl_2_, 2mM MgCl_2_, 30mM glucose, 10mM HEPES, pH = 7.35).

Halo ligand, JF646, was used at a final concentration of 200nM. Cells were incubated with JF646 for 1 hour, rinsed, and then incubated for 30 minutes (all in culture medium) before imaging in Live Cell Imaging buffer.

For FRAP experiments, 100 frames were taken every 2 seconds, followed by 20 frames every 20 seconds. 5 z planes were taken for each time frame. Photobleaching was performed after the 5^th^ time frame.

### Live cell imaging at different temperatures

To achieve rapid changes in temperature while performing live cell imaging, we used a Cherry Temp device from Cherry Biotech, and prepared cells based on the manufacturer’s manual. Briefly, COS-7 cells were seeded on 24 × 60mm No. 2 coverslips and transfected using Lipofectamine 2000 as described above. A PYREX 20 × 10 cloning cylinder was placed on the center of the coverslip using silicon grease to limit the area of transfection and was removed after transfection. Imaging was performed on LSM 880 or LSM 800 as described above 16-36 hrs after transfection. The change in the temperature occured withing 5 seconds, and the focus of the scope was adjusted manually after that. This re-equilibration of the focus and readjustment required between 2-3 minutes, which is the interval between timepoints shown in Figure 3H.

### Image processing, Analysis, and Statistics

For presentation purposes, brightness and contrasts of images were adjusted using the ImageJ software. Some of the high magnification fields were enlarged using the Adjust Size function on ImageJ.

For FRAP analysis, an average of 5 z-planes per time point was used to quantify fluorescence intensity changes and the mean fluorescence intensity of the photobleached structure was measured over time. The normalized intensity for each time-point, *I*_*normalized*_*(t)*, was calculated using the following equation

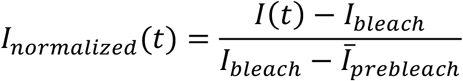

Where *I(t)* is the raw mean intensity for the FRAPed area at each time point, I_bleach_ is the mean intensity immediately after photobleaching, and 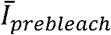 is the average intensity of the five timepoints before photobleaching. To measure raw mean intensities, the FRAPed area was tracked over time using an inhouse imageJ macro.

To quantify changes in synaptopodin-ER assemblies in response to changes in temperature, we calculated an approximate partition coefficient (PC) value. Since the small size of the synaptopodin-ER assemblies on which the analysis was performed (Figure 2G) made it difficult tracking them over time, we used an inhouse imageJ macro to automatically detect the top 100 brightest spots per time frame. We then approximated the partition coefficient (PC) for the 100 spots, as the ratio between their mean intensity to the mean intensity of the background as defined by a circle with the diameter of 3.58µm concentric to the spot.

Plots were generated using MATLAB. Protein networks were generated using R.

### Electron microscopy of brain tissue

All experiments were carried out in accordance with National Institutes of Health (NIH) guidelines and approved by the Yale IACUC. All mice were maintained in the vivarium with a 12-h light–dark cycle, stable temperature at 22 ± 1 °C and humidity between 20 and 50%. Three month old wild-type (C57BL/6) and synaptopodin KO mice were anesthetized with a ketamin/xylazine anesthetic cocktail and transcardially perfused with 2% PFA and 2% Glutaraldehyde in 0.1 M Sodium Cacodylate buffer, pH 7.4 at 37°C. Brains were dissected out, kept overnight in the same fixative and subsequently trimmed in small blocks (less than 0.5 × 0.5 × 0.5 mm), kept in fresh 2.5% Glutaraldehyde in 0.1 M Sodium Cacodylate buffer for another hour at room temperature (RT), post-fixed in 2% OsO4 + 1.5% K4Fe(CN)6 (Sigma-Aldrich) in 0.1M sodium cacodylate buffer for 1h, enbloc stained in 2% aqueous uranyl acetate for 1h, dehydrated in increasing concentrations of ethanol and embedded in EMbed 812. Ultrathin sections (50-60 nm) were observed in a Talos L 120C TEM microscope at 80 kV. Images were taken with Velox software and a 4k × 4K Ceta CMOS Camera (Thermo Fisher Scientific). Except when noted, all EM reagents were from EMS, Hatfield, PA.

**Table S2.** List of antibodies used in this study.

| Protein (epitope) | Company; Catalog number | Antibody species | Working dilution for immunocytochemistry | Working dilution for immunoblotting |
| --- | --- | --- | --- | --- |
| Synaptopodin | Sigma S9442-200UL | Rabbit | 1:1000 | 1:2500 |
| BioID2 | BioFront Tech BID2-CP-100 | Chicken | 1:1000 | NA |
| Streptavidin-Alexa Fluor 647 | Thermofisher S21374 |  | 1:1000 | 1:5000 |
| HA | Sigma 12158167001 | Rat | 1:500 | NA |
| Phalloidin-Alexa Fluor 488 | Thermofisher A12379 |  | 1:300 | NA |

**Table S3.**
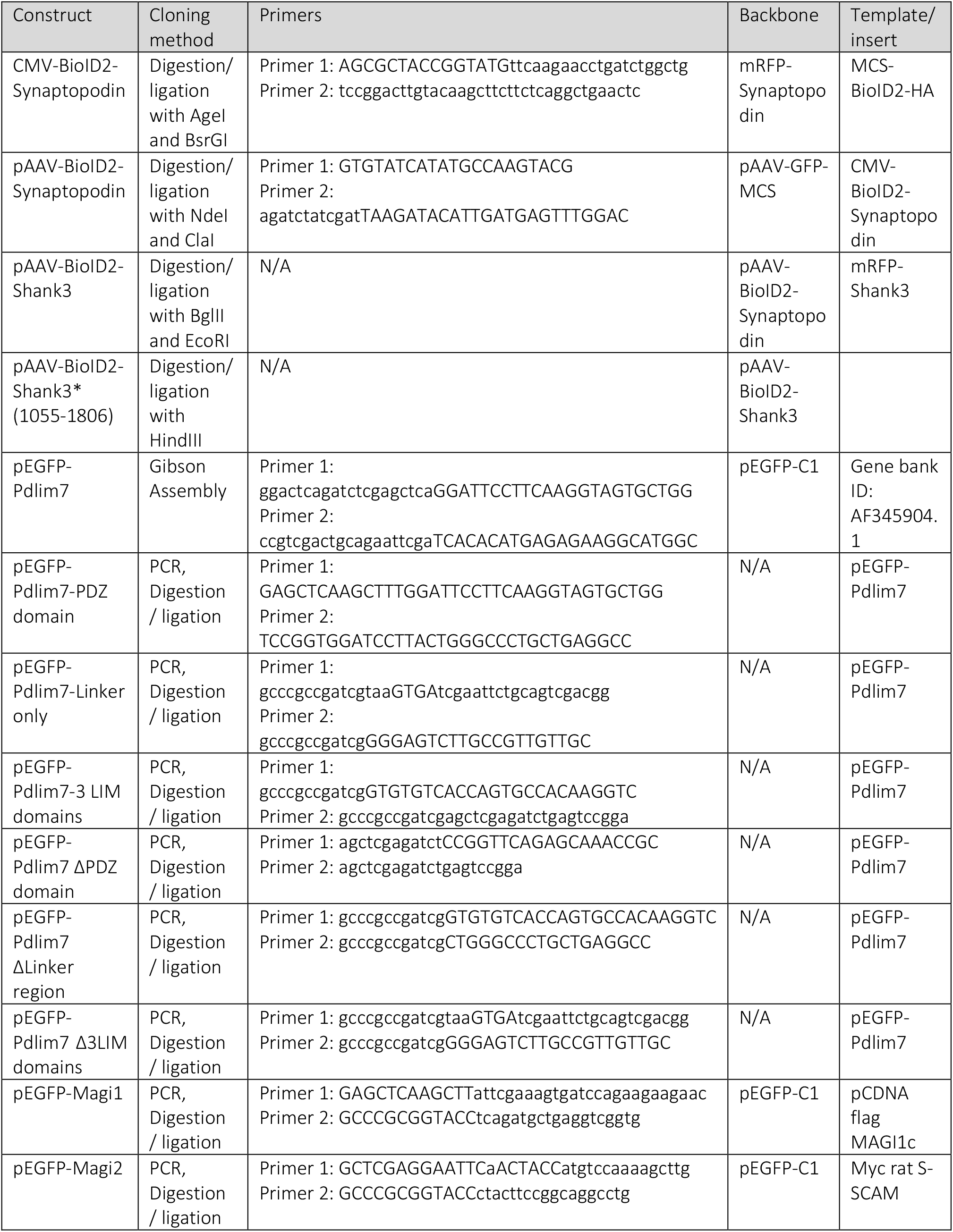
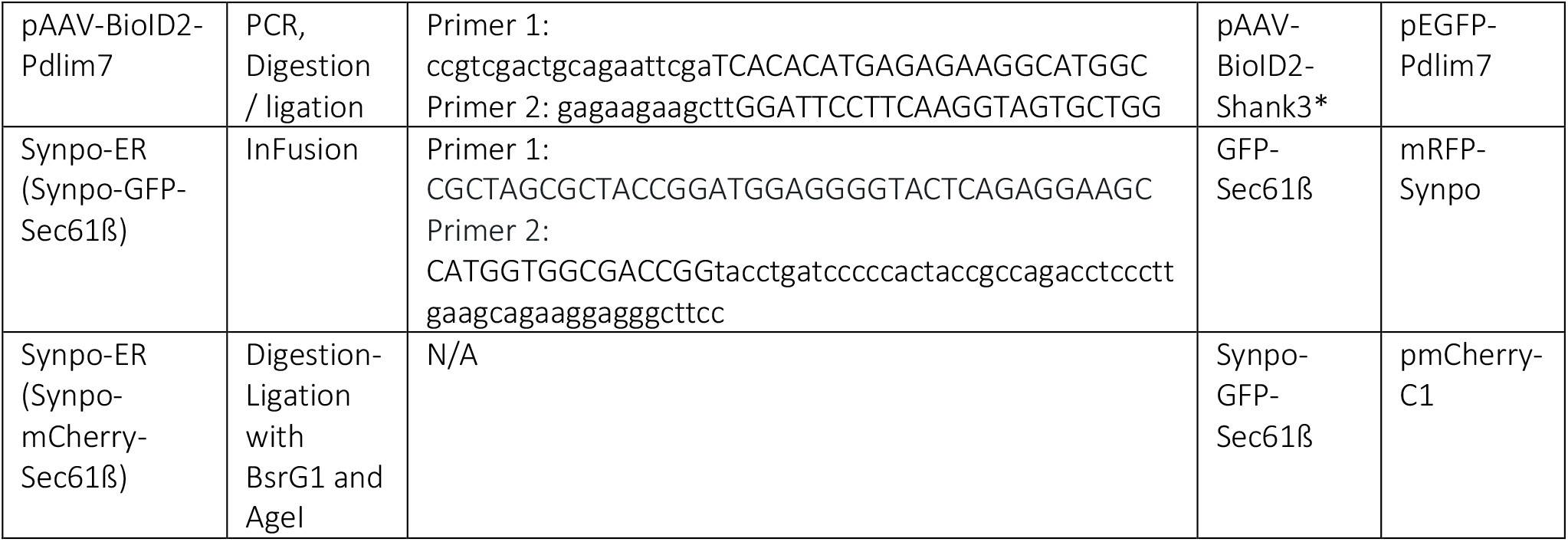

### VIDEOS

**Video 1**. Part of a dendrite reconstructed using an in house semiautomatic algorithm from FIB-SEM images of mouse cortical neurons reported previously (Hayworth et al., 2015). Plasma membrane is shown in blue, post-synaptic density in yellow, and ER in red.

**Video 2**. COS-7 cell expressing EGFP-synaptopodin (magenta) and the ER marker mCherry-VAPB (green). Treatment with 2µM Latrunculin A (LatA) resulted in the collapse of synaptopodin onto the ER. Time in min:sec.

**Video 3**. COS-7 cell expressing EGFP-VAPB (green) and mRFP-synaptopodin (magenta) and exposed to drastic hypotonic conditions. As ER vesiculates, synaptopodin clusters appeared to remain associated the ER vesicles. Z-projection is shown. Time in min:sec.

**Video 4**. COS-7 cell expressing EGFP-Pdlim7 (magenta) and the ER marker mCherry-VAPB (green). Treatment with 2µM Latrunculin A (LatA) resulted in the collapse of Pdlim7 onto the ER. Time in min:sec.

**Video 5**. COS-7 cell expressing EGFP-Pdlim7 (green) and the mRFP-synaptopodin (red) treated with 2µM Latrunculin A (LatA). Pdlim7 is associated with both latA sensitive and LatA insensitive synaptopodin assemblies. Time in min:sec.

### EXCEL TABLE

**Excel Table 1**. Relative abundance of proteins (abundance in BioID2-synaptopodin/ abundance in BioID2-Shank3*) significantly enriched in the proximity proteome of either synaptopodin or Shank3*, and p-Value of the enrichment assessed by Student’s t-test.

