## Supplementary Figures for "Properties and proximity proteomics of synaptopodin provide insight into the molecular organization of the spine apparatus of dendritic spines"

Figure S1.

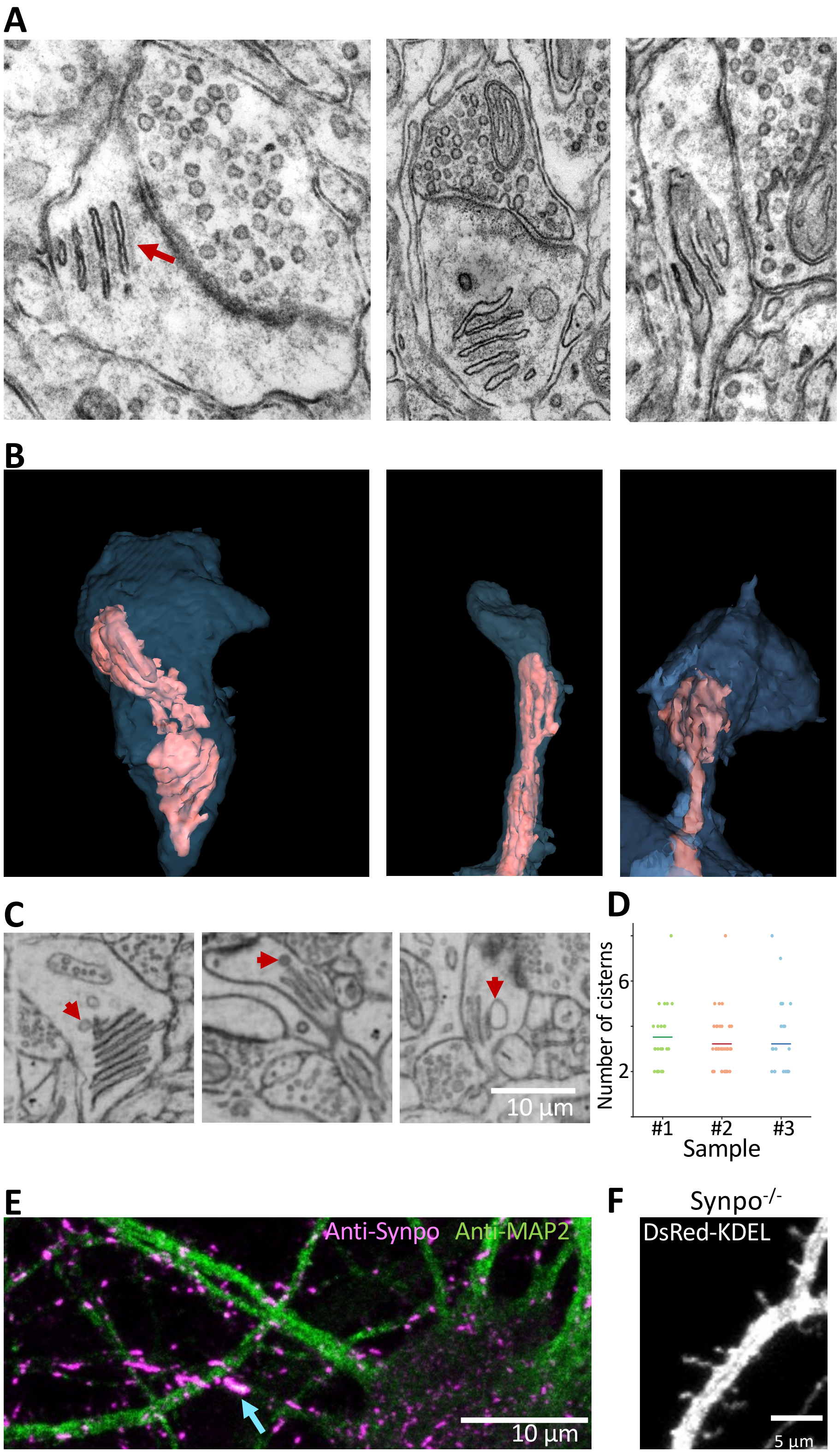

**Figure S1.** Morphological features of the SA in neurons of the mouse cerebral cortex. **A.** Examples of SA as visualized by transmission EM. Red arrow shows the dense matrix at the free surface of an outer cistern. **B.** Examples of semiautomatically reconstructed dendritic spines and SAs from 3D volumes acquired by FIB-SEM. The ER is in red and the plasma membrane in blue. **C.** Examples of SAs in contact with tubulovescicular structures (red arrows). **D.** Number of cisterns per SA in three distinct samples of cerebral cortices. Each spot represents a single SA, and the mean number of stacks for each sample is shown as a line. Data and images reported in B-D were extracted from FIB-SEM acquired 3D volumes published previously (Hayworth et al., 2015). **E.** In a subset of cultured hippocampal neurons, anti-synaptopodin staining shows the localization of this protein at axonal initial segment (blue arrow, note the lack of MAP2 staining). **F.** Presence of ER in dendritic spines of a cultured hippocampal neuron of a synaptopodin KO mouse expressing the ER marker DsRed-KDEL.

Figure S2.

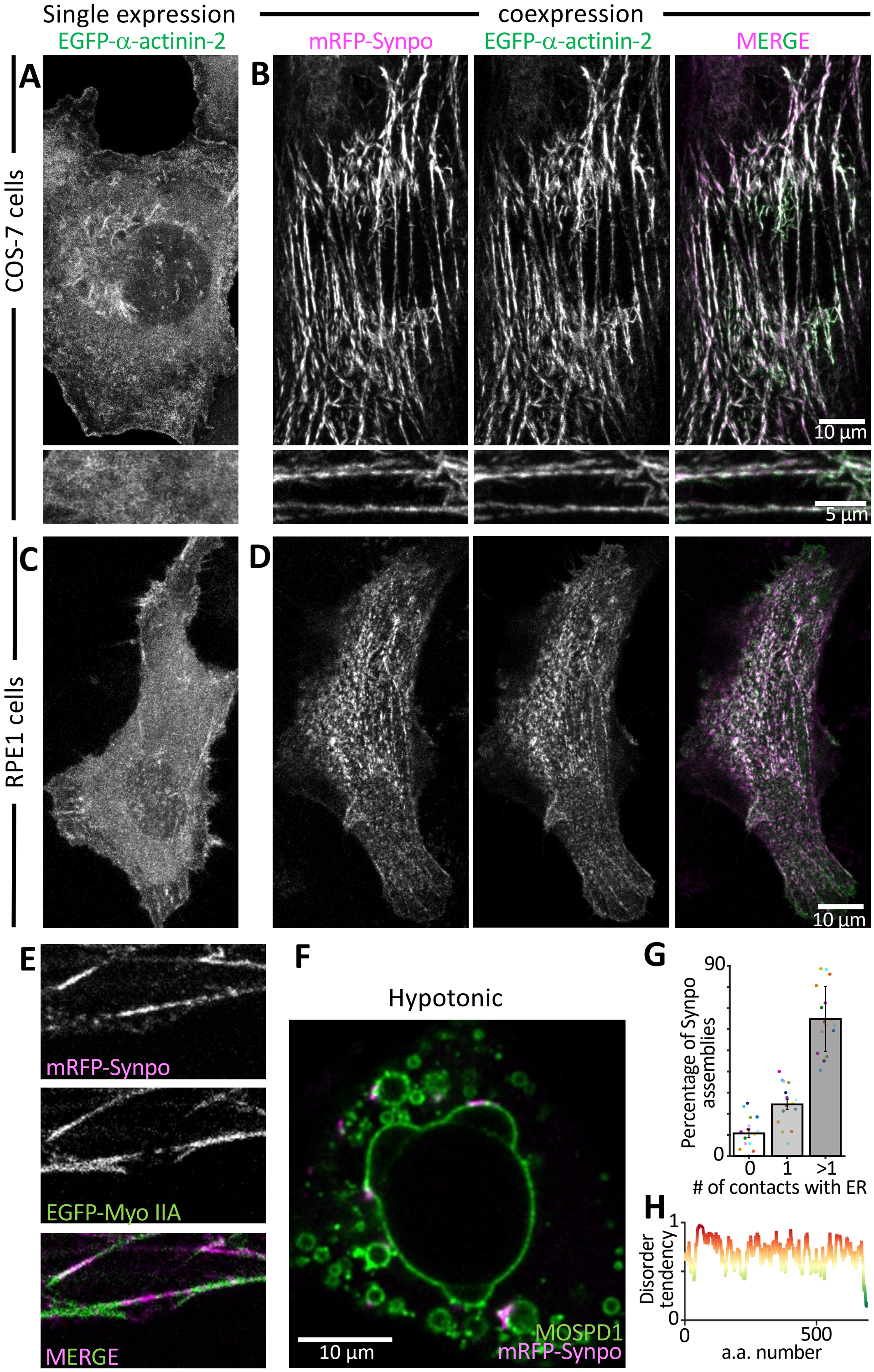
Single expression
coexpression
EGFP-α-actinin-2
mRFP-Synpo
EGFP-α-actinin-2
MERGE
A
B
COS-7 cells
10 µm
5 µm
C
D
RPE1 cells
10 µm
E
mRFP-Synpo
EGFP-Myo IIA
MERGE
F
Hypotonic
10 µm
MOSPD1
mRFP-Synpo
G
Percentage of Synpo assemblies
90
0
0
1
>1
### of contacts with ER
H
Disorder tendency
1
0
0
500
a.a. number

**Figure S2.** Relation of synaptopodin to actin. **A**-**D** Interaction of synaptopodin with $\alpha$-actinin-2. EGFP-$\alpha$-actinin-2 is cytosolic when expressed alone in COS-7 cells (**A**) or RPE1 cells (**B**), but upon coexpression with mRFP-synaptopodin $\alpha$-actinin-2 colocalizes closely with synaptopodin (**B** and **D**). **E.** Stress fiber-like assemblies of mRFP-synaptopodin alternate with Myosin IIA positive stretches in COS-7 cells. **F.** Exposing COS-7 cells to drastic hypotonic conditions causes vesiculation of the ER (labeled in this figure by EGFP-MOSPD1) and reorganization of synaptopodin assemblies that associate with ER vesicles. **G.** Quantification of the number of mRFP-synaptopodin assemblies in contact with one or more ER vesicles after hypotonic shock. Each point represents a single cell. **H.** The predicted disorder tendency for the amino acid sequence of synaptopodin as predicted by IUPred (https://iupred2a.elte.hu).

### Figure S3.

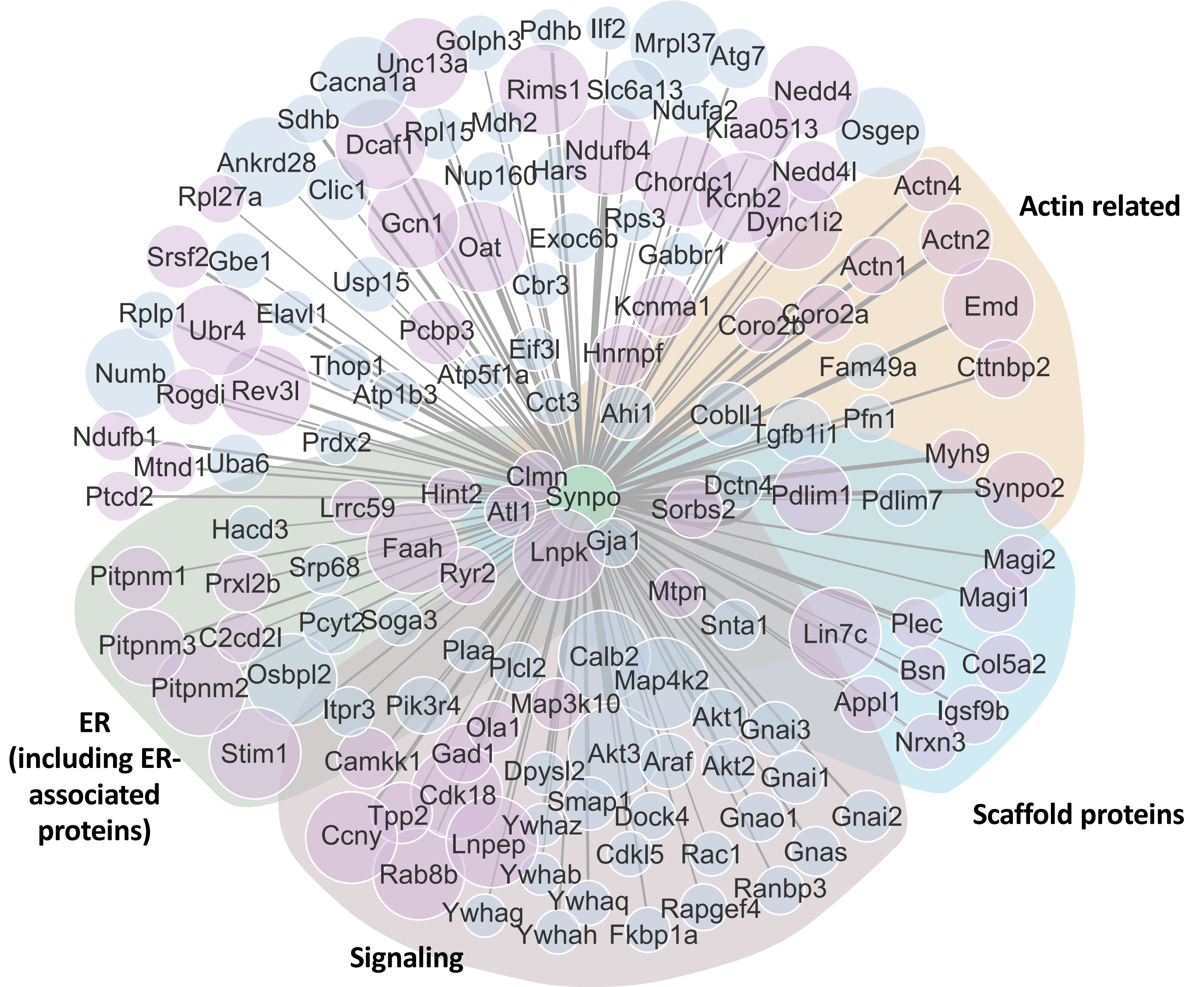


**Figure S3.** Proteins identified as neighbors of synaptopodin by proximity labeling. Proteins identified in the first and second round of proximity biotinylation are shown in blue and purple circles, respectively. The size of the circles correlates with the enrichment in BioID2-Synpo samples relative to BioID2-Shank3* samples, and the size of the lines correlates with the significance of the enrichment [-log(p_value)]. Proteins of different functional classes are grouped in different shades.

Figure S4.

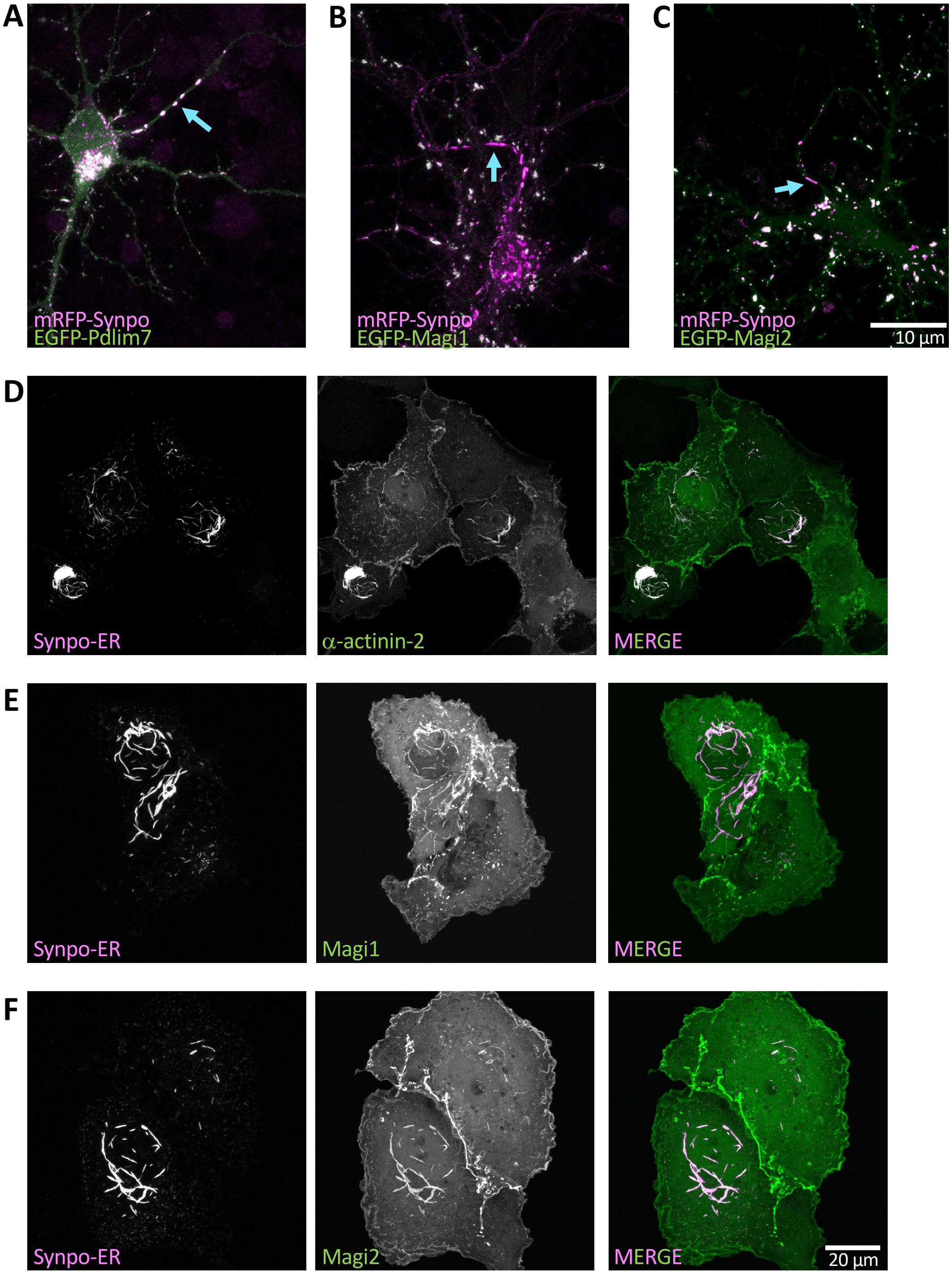


**Figure S4.** Localization of SA associated proteins identified by our screen in neurons and COS-7 cells. **A.** Pdlim7 colocalizes with synaptopodin at axonal initial segment (blue arrow). Magi1 (**B**) and Magi2 (**C**) do not colocalize with synaptopodin at axonal initial segment (blue arrow). **D-F**.

Synaptopodin interacts with α-actinin-2, MAGI1 and MAGI2 in COS-7 cells. In these cells, coexpression of synaptopodin-ER (Synpo-mCherry-Sec61ß) with EGFP-α-actinin-2 (**D**), EGFP-Magi1 (**E**) or EGFP-Magi2 (**F**) results in the recruitment of these proteins to assemblies of the ER-tethered synaptopodin.

### Figure S5.

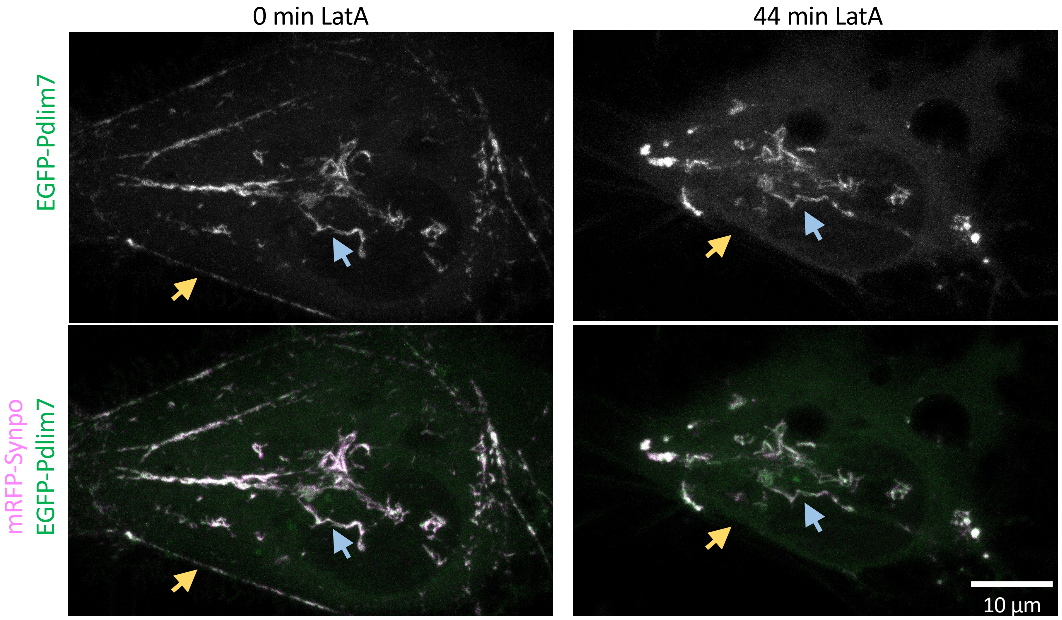


**Figure S5.** Pdlim7 colocalizes both with LatA sensitive (orange arrows) and with LatA insensitive (blue arrow) synaptopodin assemblies. COS-7 cells coexpressing EGFP-Pdlim7 and mRFP-synaptopodin were treated with 2µM LatA.

Figure S6.

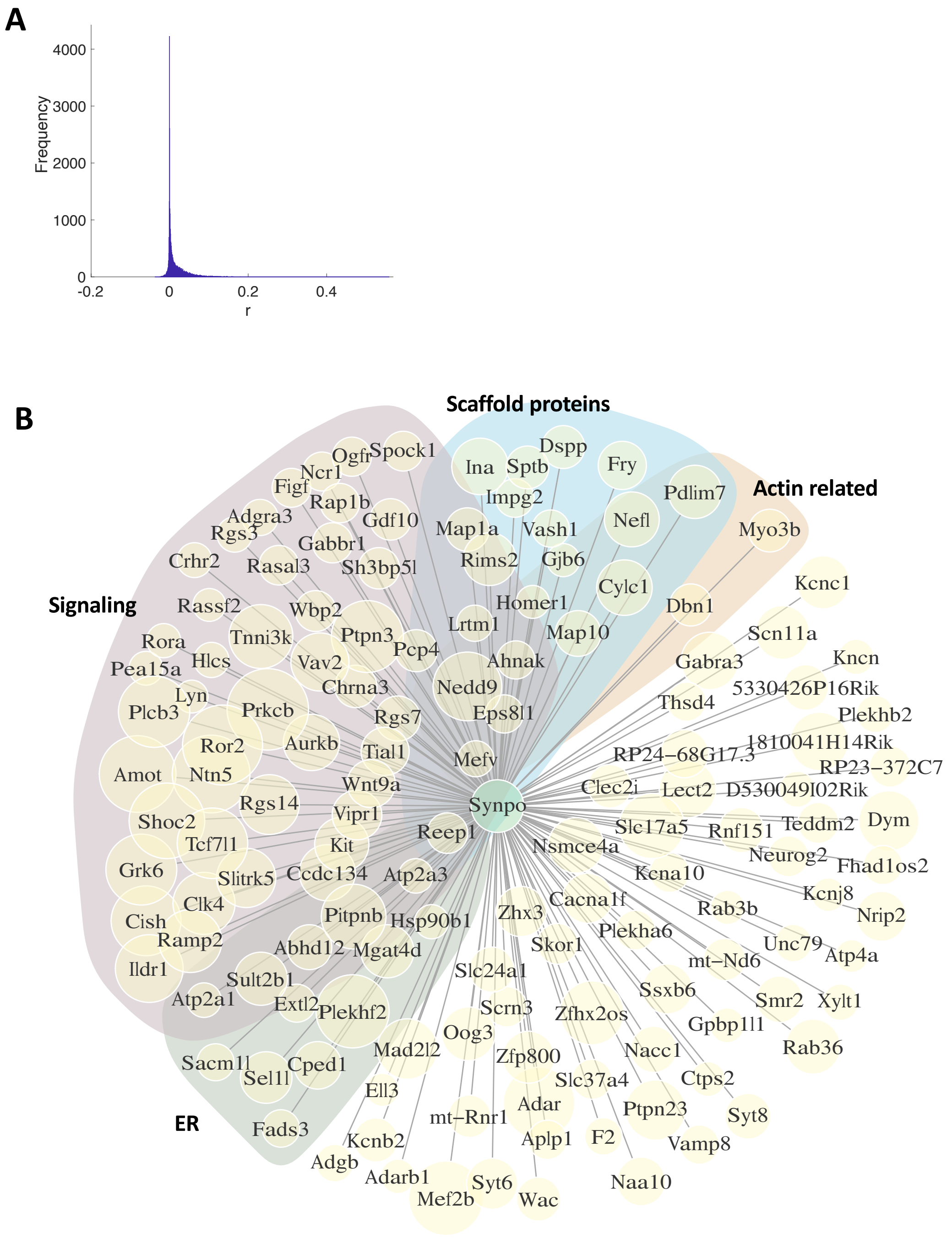


**Figure S6.** Proteins with a regional expression pattern similar to that of synaptopodin in brain. **A.** The distribution of the Pearson's correlation coefficients for the expression pattern of different proteins compared to synaptopodin is shown. **B.** Proteins with a regional expression pattern similar to that of synaptopodin in brain are depicted. These proteins have a Pearson's correlation coefficient greater than 0.2 for their expression level compared to synaptopodin throughout mouse brain. Proteins of different functional classes are grouped in different shades.
